# Regulatory T cell-derived enkephalin gates nociception

**DOI:** 10.1101/2024.05.11.593442

**Authors:** Élora Midavaine, Beatriz C. Moraes, Jorge Benitez, Sian R. Rodriguez, Joao M. Braz, Nathan P. Kochhar, Walter L. Eckalbar, Ana I. Domingos, John E. Pintar, Allan I. Basbaum, Sakeen W. Kashem

**Author notes:** **Correspondence**: Allan I. Basbaum, Chair and Professor, Department of Anatomy University of California San Francisco 1550 Rock Hall, Room 345A, San Francisco, California, USA 94158 **Lead correspondence**: Sakeen W. Kashem Assistant Professor, Dermatology University of California San Francisco 1701 Divisadero, 3rd Floor, San Francisco, CA 94115. These authors contributed equally (Allan I. Basbaum, Sakeen W. Kashem).

## Abstract

T cells have emerged as sex-dependent orchestrators of pain chronification but the sexually dimorphic mechanisms by which T cells control pain sensitivity is not resolved. Here, we demonstrate an influence of regulatory T cells (Tregs) on pain processing that is distinct from their canonical functions of immune regulation and tissue repair. Specifically, meningeal Tregs (mTregs) express the endogenous opioid, enkephalin, and mTreg-derived enkephalin exerts an antinociceptive action through a presynaptic opioid receptor signaling mechanism that is dispensable for immunosuppression. We demonstrate that mTregs are both necessary and sufficient to suppress mechanical pain sensitivity in female, but not male, mice, with this modulation reliant on sex hormones. These results uncover a fundamental sex-specific, and immunologically- derived endogenous opioid circuit for nociceptive regulation with critical implications for pain biology.

**Highlights:** 1. Gating of allodynia by meningeal Tregs is sex hormone-dependent

3. Treg-derived enkephalin modulates mechanical pain sensitivity, not inflammation

4. Delta opioid receptor on MrgprD^+^ sensory neuron mediates pain processing by mTregs

## Introduction

Pain prevalence is significantly higher in women across multiple pain conditions including nerve injury-induced neuropathic pain, musculoskeletal pain, fibromyalgia and migraine^1^. Gender disparities in pain are further evidenced by notable changes in chronic pain severity during hormonal gender affirming care^2^. Here, we identified a previously unknown immunological mechanism that underlies sex differences in both acute and chronic pain regulation.

Regulatory T cells (Tregs) are a subset of CD4^+^ T cells characterized by their immunosuppressive function and the expression of the X-linked master regulatory transcriptional factor *Foxp3*. In addition to their critical function in limiting inflammation, Tregs are also major contributors to wound healing, they regulate stem cell turnover, maintain metabolic homeostasis, facilitate placental implantation and promote maternal- fetal tolerance^8–16^. In the context of nervous system injury, Tregs mitigate pro- inflammatory cytokine interferon- (IFN-)-driven mechanical pain hypersensitivity, suppress microglia-driven nociceptive processing and can reduce astrogliosis. Additionally, they contribute to improving remyelination, thereby promoting tissue repair^17–20^. However, whether and how Tregs can directly alter neuronal activity is still unknown.

Here, we demonstrate that meningeal regulatory T cells (mTregs) are essential contributors to baseline mechanical sensitivity, and to the inhibition of mechanical pain hypersensitivity (allodynia) after nerve injury, but only in female mice. Using a well- established spared-nerve injury (SNI) model of neuropathic pain, we further show that expanding mTregs can reduce nociceptive processing independently of Treg tissue repair programs. We find that mTregs produce the endogenous opioid met-enkephalin, with female mice exhibiting increased numbers of enkephalinergic mTregs. mTreg- derived enkephalin is required for the suppression of mechanical pain hypersensitivity, but not inflammation. This distinction reveals a novel Treg function that differs from their well-established roles in immunological restraint and tissue repair.

The involvement of distinct opioid receptors in the peripheral (PNS) and central nervous systems (CNS) in the regulation of different pain modalities is well documented. Delta (δOR) and mu (μOR) opioid receptors are differentially expressed among neuronal subsets and regulate mechanical or thermal pain processing, respectively^21–23^. Enkephalin can bind to both δOR and μOR, but preferentially engages the δOR. Here we demonstrate that the anti-allodynic effects derived from mTregs are mediated by the δOR that is expressed on the *Mas*-related G protein-coupled receptor member D (MrgprD) subset of unmyelinated primary sensory neurons.

Given the pronounced sex differences observed in Treg-mediated suppression of nociceptive processing, we further explored the determinants of this sex selectivity. Our findings indicate that gonadal hormones, rather than sex chromosomes, play a key role in modulating Treg suppression of nociceptive thresholds. Thus, we propose a novel mechanism by which Tregs mediate the suppression of nociception, regulated by sex- specific hormonal influences.

## Results

### Sex-specific suppression of nociceptive thresholds by meningeal Tregs

As the immune system has emerged as a central determinant driving sex differences in pain sensitivity, we sought to investigate its contribution to nociceptive processing^24^. Tregs are a fundamental cell type essential for maintaining and restoring tissue homeostasis. Here we initially focused on identifying Treg localization and function within nervous system tissues. Using confocal microscopy and flow cytometry on whole meningeal sheets, dorsal root ganglia (DRG), sciatic nerves, spinal cords (SC) and brain, we localized Tregs to the meninges of the CNS, and to the leptomeninges of the DRG (**Figure 1A-C**). To enhance the sensitivity of Treg detection within these organs, we utilized the bright reporter signals from the double reporter *Foxp3*^eGFP-Cre-ERT2^*;Rosa26^LSL-tdTomato^* mice. Notably, and consistent with previous reports we observed a more pronounced localization of Tregs in the lumbar and caudal segments of the spinal cord meninges (**Figure 1A**) ^25^. In the DRG, as for most leukocytes other than resident macrophages, Tregs predominated in the leptomeninges, proximal to the dorsal roots entry zone, with sparse presence within the DRG parenchyma (**Figure 1B**)^26^. By intravenous administration of a fluorescently-labeled CD45 antibody and using flow cytometry, we distinguished intravascular Tregs from tissue Tregs **(Figure S1A)**^27^. We quantified the numbers of non-vascular, tissue Tregs in various organs within the nervous and lymphoid systems. We will further refer to the SC meningeal and DRG leptomeningeal Tregs as mTregs (**Figure 1C**). We observed minimal localization of Tregs in peripheral nerves and did not detect any Tregs within the parenchyma of the CNS in young, uninjured mice. We observed nearly equivalent numbers of tissue Tregs between male and female mice across tissue (**Figure 1D**)^18^.

**Figure 1.**
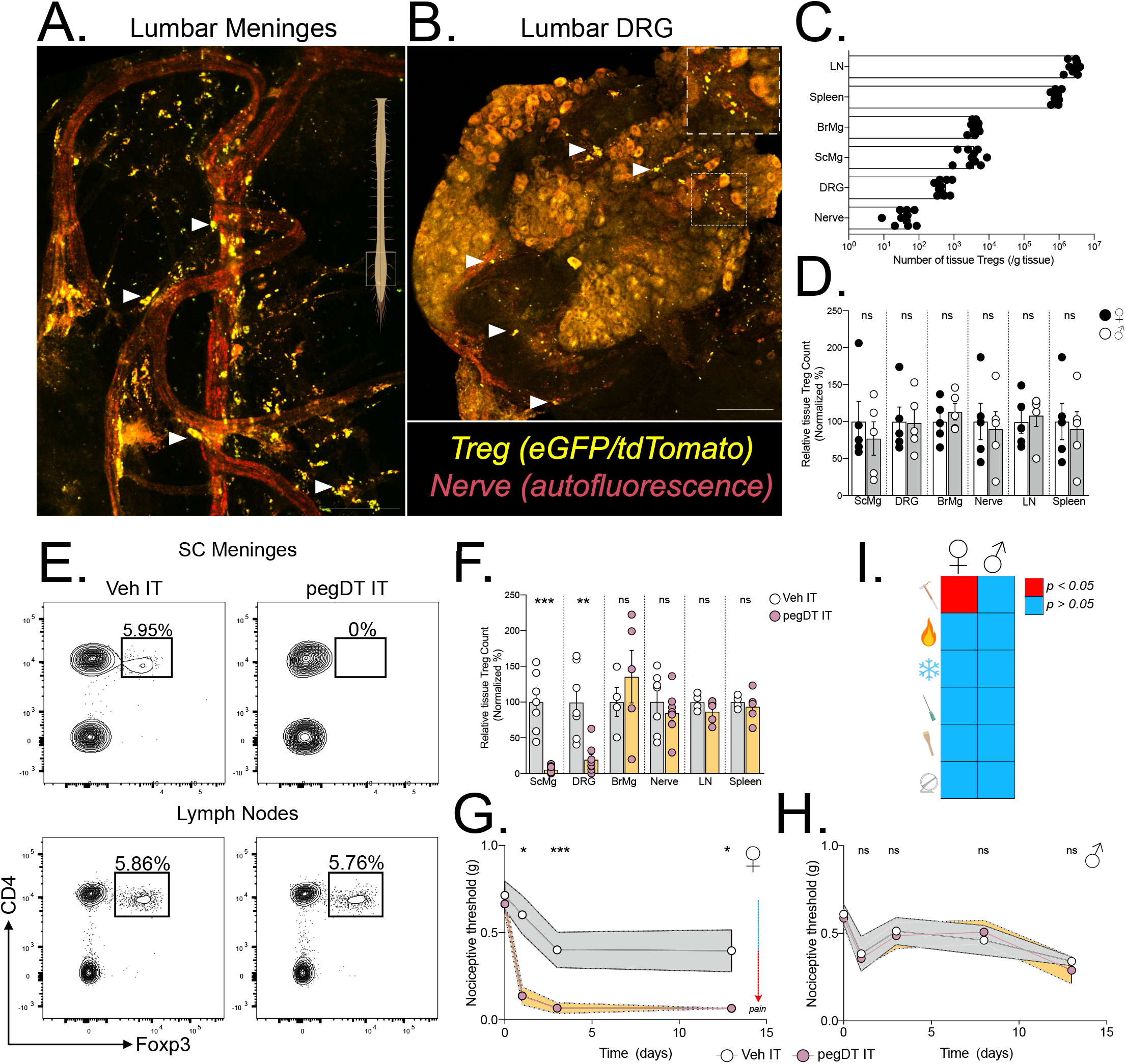
mTreg suppress mechanical pain hypersensitivity in female mice. Representative whole mount maximum projection confocal microscopy image of the (A) lumbar spinal cord meninges and (B) DRG showing Tregs (green-red: yellow) and nerves (autofluorescence, red) in *Foxp3*^eGFP-CreERT2^*;Rosa26*^tdTomato^ reporter mice. Inlet showing DRG magnification. Scale bar represents 100 µm in A) and 150 µm in B). Arrows indicate Tregs. (C) Total number of weight-adjusted tissue Tregs across organs, in both sexes combined. (D) Relative number of tissue Tregs from male (white) and female (black) mice per organ. 100% represents mean number of female Tregs per organ. Comparison is made between each individual organ. (E) Representative concatenated flow cytometry plots of tissue Treg after a single intrathecal (IT) injection of 20 ng of pegylated diphtheria toxin (pegDT). (F) Relative quantifications of tissue Treg depletion 2 days after a single IT pegDT injection across organs. 100% represents mean number of tissue Tregs in IT vehicle-injected mice per organ. (G and H) 50% paw withdrawal thresholds measured using von Frey filaments before (day 0) and after a single dose of 20 ng of IT pegDT or vehicle in female (G) or male (H) *Foxp3*-DTR mice. (I) Summary of significant behavioral differences comparing IT pegDT- and control-injected female and male mice. Total number of mice for G-I is presented in Figure S3. ScMg= Spinal cord meninges, BrMg= Brain meninges, LN= Lymph nodes, ns = not significant, \**p*<0.05, \*\**p*<0.01, \*\*\**p*<0.001. **Related to Figure S1-S3.**

To assess the feasibility of site-specific depletion of mTregs, we performed intrathecal (IT) injections of pegylated diphtheria toxin (pegDT) in *Foxp3-*DTR mice^28^. Although an IT injection of Evan’s blue rapidly spreads through the SC meninges and DRG, into the brain and to the draining cervical and lumbar lymph nodes, pegylated fluorescently labeled molecules (pegDyLight 650) remain restricted to the SC meninges and to the DRG **(Figure S2A-B)**. Consistently, a single 20 ng dose of pegDT IT selectively depleted >90% of SC and DRG mTregs in both male and female mice, but spared Tregs located in the brain meninges, draining lymph nodes, spleen, and peripheral nerves (**Figure 1E-F**). Importantly, *Foxp3*-DTR mice subjected to repeated IT administrations of pegDT do not exhibit the significant weight loss, splenomegaly and mortality that typically develops following systemic autoimmunity in *Foxp3*-DTR mice induced by repeated intraperitoneal (IP) injections of diphtheria toxin (DT) **(Figure S2C- F)**. Clearly, the pegDT IT system offers a novel method for selective depletion of mTregs while avoiding systemic inflammation.

We next evaluated behavioral outcomes in mice following mTreg depletion. A single dose of pegDT IT induced a profound and prolonged decrease in mechanical thresholds in naïve female but not male *Foxp3*-DTR mice (**Figure 1G-H**). Importantly, mechanical thresholds in wildtype (WT) C57BL/6 mice treated with pegDT IT or *Foxp3*-DTR mice treated with vehicle IT did not differ, ruling out pegDT or IT injections as the cause of the sex-specific allodynia **(Figure S3A)**. In addition to evaluating mechanical hypersensitivity, which is conveyed by mechanosensitive unmyelinated and myelinated primary afferent nerve fibers, we also assessed mice for noxious heat sensitivity mediated by Trpv1^+^ nociceptors, cold sensitivity that is mediated by Trpm8^+^ nociceptors, pin prick sensitivity mediated by Aδ afferents and brush responses mediated by Aβ fibers. Although depletion of mTregs selectively induced mechanical allodynia in females, it did not impact any other sensory modality. Motor function tests using the rotarod also did not differ in either sex (**Figure 1I and Figure S3A-H)**. We conclude that mTregs selectively suppress mechanical thresholds in a sex-dependent manner, effectively preventing mechanical allodynia in a previously uninjured state.

### Expansion of mTregs alleviates injury-induced mechanical allodynia independently of tissue repair

In addition to exploring mTreg role in mechanical sensitivity in uninjured mice, we investigated whether mTreg can suppress allodynia following nerve injury. Using a well-established spared nerve injury (SNI) model of neuropathic pain, we transected and ligated the common peroneal and tibial nerve branches of the sciatic nerve, sparing the sural nerve. This model induces chronic, unremitting, and permanent mechanical hypersensitivity with a non-healing neuroma formation four weeks after the injury (**Figure 2A-B**)^29,30^. As mice with SNI exhibit mechanical thresholds at the limit of detection with commercially available von Frey filaments, we conducted single fiber testing using the lowest available 0.008 g von Frey filament. Again, mTreg depletion increased allodynia following SNI in females, but not in males (**Figure 2C-D**).

**Figure 2.**
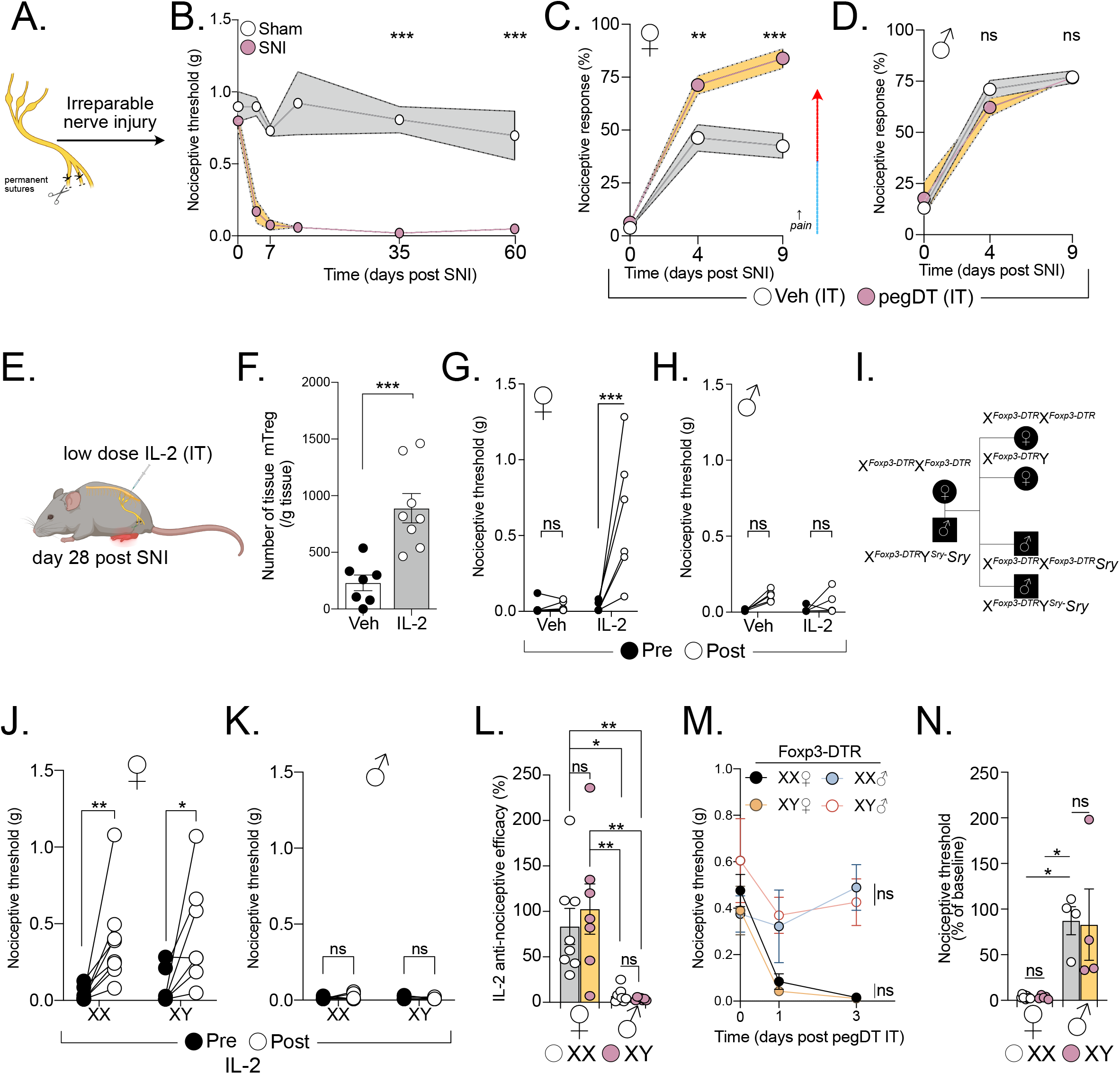
Expanding mTreg alleviates nociception dependent on sex hormones and independent of tissue repair. (A) Schematic representation of the spared nerve injury (SNI) surgery. (B) Long-term assessment of mechanical thresholds in mice following SNI surgery (both sexes combined, no difference between the sexes). n *=* 7-8 mice per group. (C and D) Percent response to 0.008 g von Frey filament in mice with SNI (day 0) and treated with IT pegDT or vehicle every 4 days. n = 8 per group for females and 9-10 per group for males. (E) Schematic representation of mTreg expansion in mice 4 weeks after SNI by 3 IT injections of low-dose IL-2 (0.1 µg). (F) Total mTreg number in meninges after low-dose IL-2 or vehicle IT injections (both sexes combined, no differences between the sexes). (G and H) Nociceptive thresholds of females (G) and male (H) mice given low-dose IL-2 or vehicle IT 4 weeks after SNI. (I) Schematic representation of the mating strategy of Four Core Genotypes (FCG) *FoxP3*-DTR mice demonstrating resulting XX and XY females and XX and XY male mice. (J and K) Nociceptive thresholds of FCG female (J) and male (K) mice following low-dose IL-2 or vehicle IT injections 4 weeks after SNI. (L) Anti-nociceptive efficacy determined as post IL-2/vehicle injection threshold divided by baseline mechanical threshold in male and female mice with XX (white) or XY (pink) chromosomes. (M) Nociceptive thresholds of FCG *FoxP3*-DTR female and male mice following a single IT pegDT or vehicle injection. (N) Percent baseline nociceptive thresholds determined as post pegDT/vehicle injection threshold divided by baseline mechanical threshold in male and female mice with XX (white) or XY (pink) chromosomes. ns = not significant, \**p*<0.05, \*\**p*<0.01,\*\*\**p*<0.001. **Related to Figure S4.**

We next asked whether expanding mTregs could alleviate the mechanical allodynia independently of tissue repair. Tregs express the high affinity interleukin-2 receptor, IL- 2Rα, and low-doses of IL-2 can effectively expand Tregs in mice, a therapeutic approach that has been used to treat autoimmune diseases in humans^31^. IT injections of low-dose IL-2 successfully expanded mTregs in both male and female mice (**Figure 2F**). However, although mTreg expansion promoted significant anti-allodynia in SNI female mice, it did not exhibit a similar effect in males (**Figure 2G-H**). It is noteworthy that acute IT injection of IL-2, in uninjured mice, did not increase nociceptive thresholds, suggesting that an IL-2 or Treg-based therapy could selectively improve neuropathic pain without affecting basal nociceptive processing **(Figure S4A-B)**. IL-2 injections and mTreg expansion in mice with SNI likewise did not alter noxious cold or heat sensitivity, again highlighting the specificity of the sensory modality modulation by mTreg in the context of neuropathic pain **(S4C-E)**.

### Gonadal hormones, not sex chromosomes determine sex-selective, anti- nociceptive function of mTregs

*Foxp3* is an X-linked gene, some of which escape X-inactivation. Moreover, random X-inactivation has been suggested to be potentially altered during inflammatory state in females^32–34^. To test whether sex chromosomes dosage contributes to our observed phenotype, we used the Four Core Genotypes (FCG) mouse model in which gonadal sex in mice is independent of sex chromosomes^35,36^. FCG mice harbor a deficiency in the sex determining region Y protein (*Sry*) on the Y chromosome and instead feature an autosomal transgenic insertion of *Sry.* This genetic configuration enables the discrimination of sex chromosome dose influence from the contribution of gonadal hormones (**Figure 2I**). Both XX and XY chromosome gonadal female mice displayed mTreg-mediated alleviation of mechanical allodynia after SNI; XX- and XY- gonadal male mice did not (**Figure 2J-L**). Similarly, after mTreg depletion in *Foxp3*-DTR mice crossed to the FCG system, we found that female specific gonadal hormones, but not sex chromosome, mediate the mTreg suppression of nociceptive thresholds in the absence of injury (**Figure 2M-N**). Based on our findings in both uninjured and chronic injury states, we conclude that there is a profound and consistent sex hormone-dependent contribution of mTregs to the modulation mechanical pain sensitivity.

### Regulatory T cells express the endogenous opioid peptide enkephalin

To investigate the molecular mechanisms though which Tregs suppress nociceptive thresholds, we first interrogated public genomic resources. We hypothesized that meningeal tissue Tregs could exhibit an activated lymphoid Treg phenotype rather than a resting Treg phenotype. Tregs have increased expression of the *Penk* gene, which encodes for *Proenkephalin*, a peptide precursor of both Met- and Leu-enkephalin, in various tissues including the nervous system, in both mice and humans^19,37–39^. Here we re-analyzed raw public RNA-seq data of activated Tregs, resting Tregs, as well as activated and resting CD4^+^ Foxp3^-^ conventional T cells (Tconv)^40^. Strikingly, in activated versus resting Tregs, we observed a significant upregulation of *Penk* expression (**Figure 3A**). We also investigated other opioid ligand and receptor genes but only recorded a very sparse expression of other opioid-related genes among the CD4^+^ T cell subsets (**Figure 3B**). Based on our prior experience defining mechanical sensitivity through enkephalin-δOR signaling^21^, we therefore focused on Treg expression of *Penk.* By ATAC-seq analysis, we observed open chromatin regions of the *Penk* locus in activated Tregs, but not in other CD4^+^ T cell subsets. This open chromatin was similar to the open chromatin, promoter and enhancer regions of the developing forebrain, an established enkephalin-producing area of the murine CNS (**Figure 3C**)^41^. By analyzing the raw dataset from the Immunological Genome Project^42^, we also explored *Penk* expression within cell populations of the immune system. We observed a strikingly greater *Penk* expression in Tregs, compared to other immune cells (**Figure 3D**). Furthermore, *Penk* expression in Tregs increase significantly following stimulation with IL-2, compared to other common gamma chain cytokines (**Figure 3E**).

**Figure 3.**
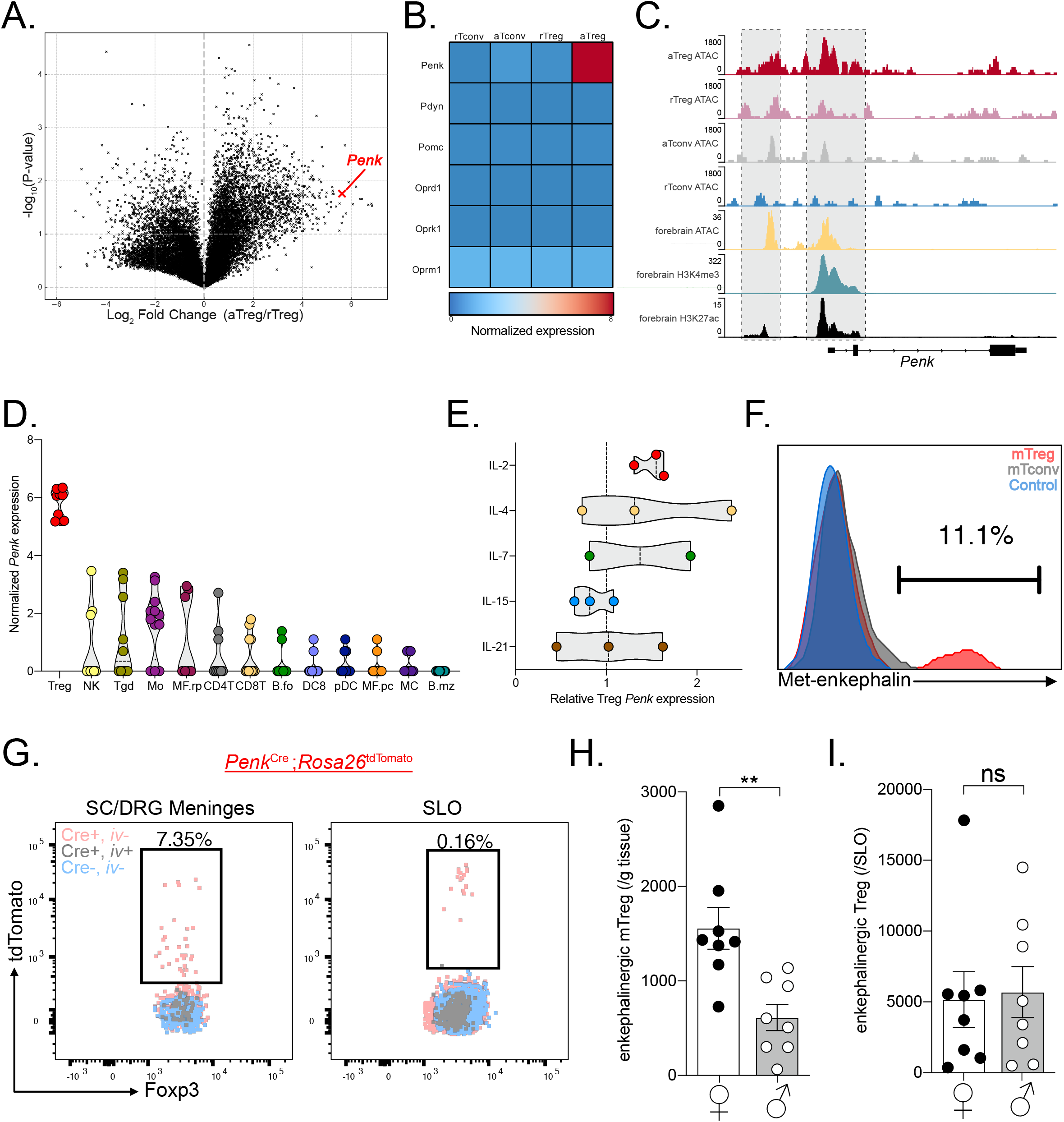
mTregs express and produce enkephalin. (A) Volcano plot of transcription fold change of activated Treg (aTreg) vs resting Treg (rTreg) and (B) heatmap of relative log2 expression value of aTreg, rTreg and activated and resting CD4^+^ CD25^-^ conventional T cells (aTconv and rTconv) from public dataset GSE154680 (n=3). (C) Averaged ATAC sequencing (ATACseq) of open chromatin accessibility peaks on the *Penk* locus in different T cell subsets (n=4 per group, GSE154680), compared to ATACseq, and histone modification Chip-Seq from public ENCODE dataset of the p0 developing forebrain, a known enkephalinergic region. (D and E) Log2 values of *Penk* expression by different unstimulated immune cell types from Immgen dataset GSE180020. (E) Treg *Penk* expression fold change after cytokine stimulation compared to vehicle control. (F) Representative PMA:Ionomycin stimulated mTregs, meningeal CD4^+^ T cells (mCD4) from WT mice or *Penk^-/-^* mTreg (control). (G) Representative flow cytometry plots of Tregs from meninges or secondary lymphoid organs (SLO) from *Penk*^Cre^*Rosa*26^tdtomato^ mice. Pink represents non-vascular, tissue Tregs from transgenic *Penk* lineage reporter mice. Gray represents vascular Treg in reporter mice while Blue corresponds to tissue Treg from non-transgenic control mice. (H-I) Number of enkephalin lineage fate reporter positive tissue Tregs in (H) meninges and (I) secondary lymphoid organs (SLO) in male and female mice. NK: Natural Killer cells, Tgd: γδ T cells, Mo: Monocytes, MF.rp: Red pulp macrophages, CD4T: CD4^+^ T cells, CD8T: CD8^+^ T cells, B.fo: splenic follicular B cells, DC8: CD8^+^ dendritic cells, pDC: splenic plasmacytoid, MF.pc: peritoneal macrophages, MC: myeloid cells, B.mz: splenic marginal zone B cells, ns = not significant, \**p*<0.05, \*\**p*<0.01,\*\*\**p*<0.001.

To establish whether Tregs indeed produce the endogenous opioid peptide enkephalin, we screened commercially available anti-Met-enkephalin antibodies. Met-enkephalin was chosen over leu-enkephalin as the latter can be cleaved from both proenkephalin and prodynorphin peptides^43^. These antibodies were validated using *Penk^-/-^* mice as negative controls. Figure 3F shows that mTregs produce met-enkephalin, but meningeal CD4^+^ T cells and lymphoid Tregs produce very low levels even after cytokine stimulation (**Figure 3F**). We validated this finding by generating *Penk*^Cre^*;Rosa26*^tdTomato^ mice, which fate-labeled enkephalinergic cells. Consistent with our antibody finding, we observed very similar number of enkephalinergic lineage (tdTomato positive) mTregs in naïve mice. Very few lymphoid or intravascular Tregs were tdTomato labeled (**Figure 3G**). Most interestingly, female mice exhibited significantly greater numbers of enkephalin-positive Tregs in the meninges, but not in the lymphoid organs. This distinction suggests that Treg fate and function variation across the sexes may be organ system specific (**Figure 3H**).

### mTreg-derived enkephalin is required for suppressing nociceptive processing

Using *Penk*^Cre^*;Rosa26*^tdTomato^ mice, we next investigated enkephalin lineage positive cells in the meninges and the DRG. Interestingly, we observed a significant increase in the representation of mTregs in the tdTomato-positive enkephalin subpopulation compared to tdTomato-negative cells (**Figure 4A-B**). In order to manipulate the enkephalin-producing immune cells, we generated bone marrow chimeric mice by transplanting *Penk*^Cre^*;Rosa26*^DTR^ bone marrow into irradiated CD45.2 congenically marked WT mice (**Figure 4C**). This strategy enables a selective DT-induced depletion of hematopoietic enkephalinergic cells, that spares depletion of non-hematopoietic enkephalinergic cells of the nervous system and the stroma. Importantly, CD4^+^ T cells of the meninges are predominantly bone marrow-derived and exhibit a tissue circulatory characteristic rather than acquiring tissue residency^44^. Consistently, we found that mTregs are indeed bone marrow-derived, similar to lymphoid Tregs, and differ from spinal microglia and skin Langerhans cells, which are host-derived **(Figure S5A-C)**. pegDT IT administrations in *Penk*DTR^Δheme^ chimeric mice decreased the number of mTregs and led to profound mechanical hypersensitivity in female, but not male mice, in both uninjured and nerve injured states (**Figure 4D-G**). We conclude that blood-derived meningeal enkephalinergic cells gate mechanical hypersensitivity in females.

**Figure 4.**
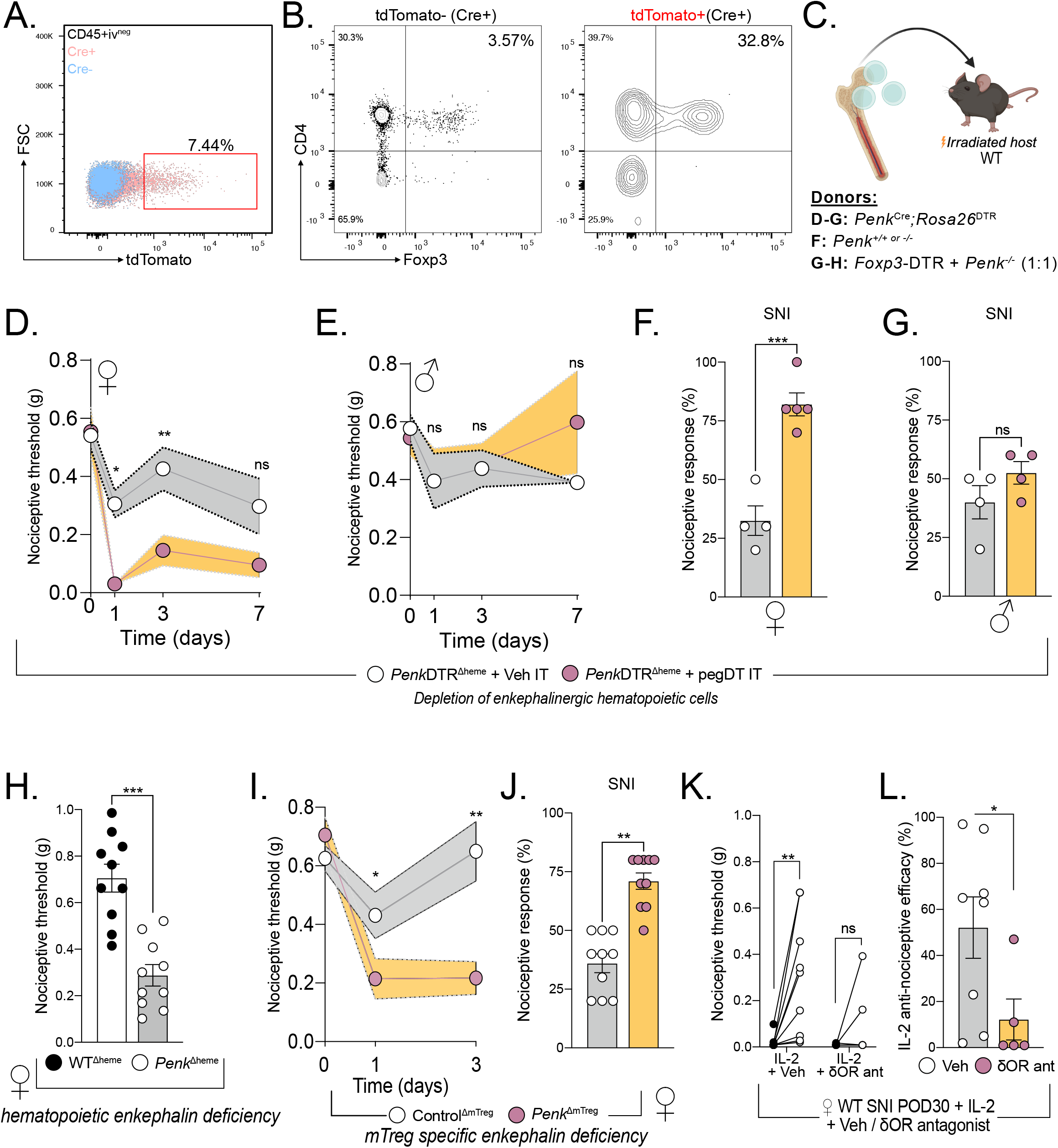
mTreg-derived enkephalin controls nociceptive thresholds. (A) Representative flow cytometry plots of Cre positive (pink) or Cre negative (blue) CD45^+^ non-vascular cells from the meninges and the DRG, combined, from *Penk*^Cre^*Rosa26*^tdTomato^ mice. (B) Flow plot shows representative tdTomato negative and right plot shows tdTomato positive leukocytes, demonstrating the more pronounced mTreg representation in the enkephalinergic fate cell population. (C) Schematic representation of bone marrow transplants to generate a global depletion of enkephalinergic cells, a depletion of hematopoietic-derived enkephalin or a depletion of Treg-derived enkephalin, respectively. (D-G) Bone marrow chimera of *Penk*^Cre^*Rosa26*^DTR^ ➔ irradiated WT recipients. Nociceptive thresholds after a single pegDT (pink) or vehicle (white) IT injection in female (D) and male (E) mice. n= 5 per group. Nociceptive thresholds after SNI and pegDT (pink) or vehicle (white) IT injection in female (F) and male (G) mice. (H) Nociceptive thresholds at baseline of female WT ➔ WT (black) or *Penk^-/-^* ➔ WT (white) bone marrow chimeras. (I) Female *Foxp3*-DTR + *Penk^-/-^*(1:1) ➔ WT mice and tested for nociceptive thresholds after pegDT (pink) or vehicle (white) IT, n=10 per group. (J) Nociceptive thresholds in SNI mice after pegDT (pink) or vehicle (white) IT injections. (K) WT SNI female mice given low-dose IL-2 and naltrindole. (L) Nociceptive efficacy calculated as percent compared to baseline threshold. ns = not significant, \**p*<0.05, \*\**p*<0.01,\*\*\**p*<0.001.

Having established a female-specific contribution of the bone marrow-derived enkephalin system, we next used female mice to dissect the mechanism of pain regulation by mTregs. To establish whether bone marrow-derived enkephalin is required for the regulation of nociceptive thresholds, we generated bone marrow chimeras in which *Penk* deficient bone marrow is transplanted into irradiated hosts, thus generating *Penk*^Δheme^ mice. As predicted, these *Penk*^Δheme^ mice display decreased nociceptive thresholds during uninjured state, compared to vehicle-injected *Penk*^Δheme^ mice, which supports our conclusion that hematopoietic cell-derived enkephalin controls basal mechanical sensitivity but only in females (**Figure 4H**).

We recognize that recombination-based selective ablation of enkephalin on Tregs, using *Foxp3^Cre^* or *Foxp3^Cre-ERT2^*has multiple limitations and caveats. These include systemic targeting of Tregs, including enkephalinergic Tregs in the skin, potential compensatory *Penk* regulation upon constitutive ablation, potential side effects of tamoxifen, potential stochastic deletion of *Penk* outside of Tregs in homozygote Cre/Cre-ERT2 mice, and the impact of random X-inactivation on heterozygous mice^37^. In light of these concerns, we also generated mixed bone marrow chimeras using 1:1 ratio of *Foxp3*-DTR and *Penk*^-/-^ bone marrow and implanted these chimeras in irradiated WT mice. Intrathecal injection of pegDT into these mice results in ablation of *FoxP3*- DTR Tregs; the remaining Tregs are left deficient for *Penk* (*Penk*^ΔmTreg^ mice) while preserving other immune cell types. Importantly, this approach circumnavigated potential depletion of any previously unrecognized non-hematopoietic cells that express *Foxp3*. At baseline, uninjected mixed chimeric mice had similar mechanical thresholds as *WT*^Δheme^ control chimeras. As predicted, pegDT IT injection led to mechanical hypersensitivity in uninjured *Penk*^ΔmTreg^ but not *WT*^Δheme^ control mice and exacerbated nerve injury-induced hypersensitivity (**Figure 4I-J**). Having shown that IL-2-induced mTreg expansion and expression of enkephalin alleviates neuropathic pain, we next investigated whether this could be mediated by the δOR, the preferred receptor for enkephalin. In these studies, we co-administered IL-2 and naltrindole, a selective antagonist of the δOR, and observed that IL-2-induced anti-allodynia was abolished (**Figure 4K-L**). We conclude that mTreg-derived enkephalin is required for suppressing mechanical pain hypersensitivity and that this suppression is mediated by the δOR.

### Treg-derived enkephalin is dispensable for immune suppression

Previously, Tregs have been shown to suppress hyperalgesia following nerve injury by suppressing IFN- -induced primary afferent sensitization^17^. Thus, we hypothesized that a potential mechanism by which Treg-derived enkephalin mediates the suppression of nociceptive thresholds involves modulation of immunological responses. To address this possibility, we tested the nociceptive thresholds of immunodeficient *Rag2^-/-^* mice which are missing both T and B cells and compared them to immunocompetent littermates. In these studies, we mated *Rag2^+/-^* mice and were surprised to observe decreased nociceptive thresholds in the *Rag2^-/-^* offspring compared to their *Rag2^+/+^* and *Rag2^+/-^* littermates. This finding suggests that there may be a mechanism of Treg-mediated control of nociceptive thresholds, which is independent of exaggerated lymphocyte-driven inflammation (**Figure 5A**). Furthermore, consistent with this conclusion, depleting macrophages through liposomal clodronate administration did not reverse the mechanical allodynia observed in female mice deficient in mTreg (**Figure 5B**)^45^.

**Figure 5.**
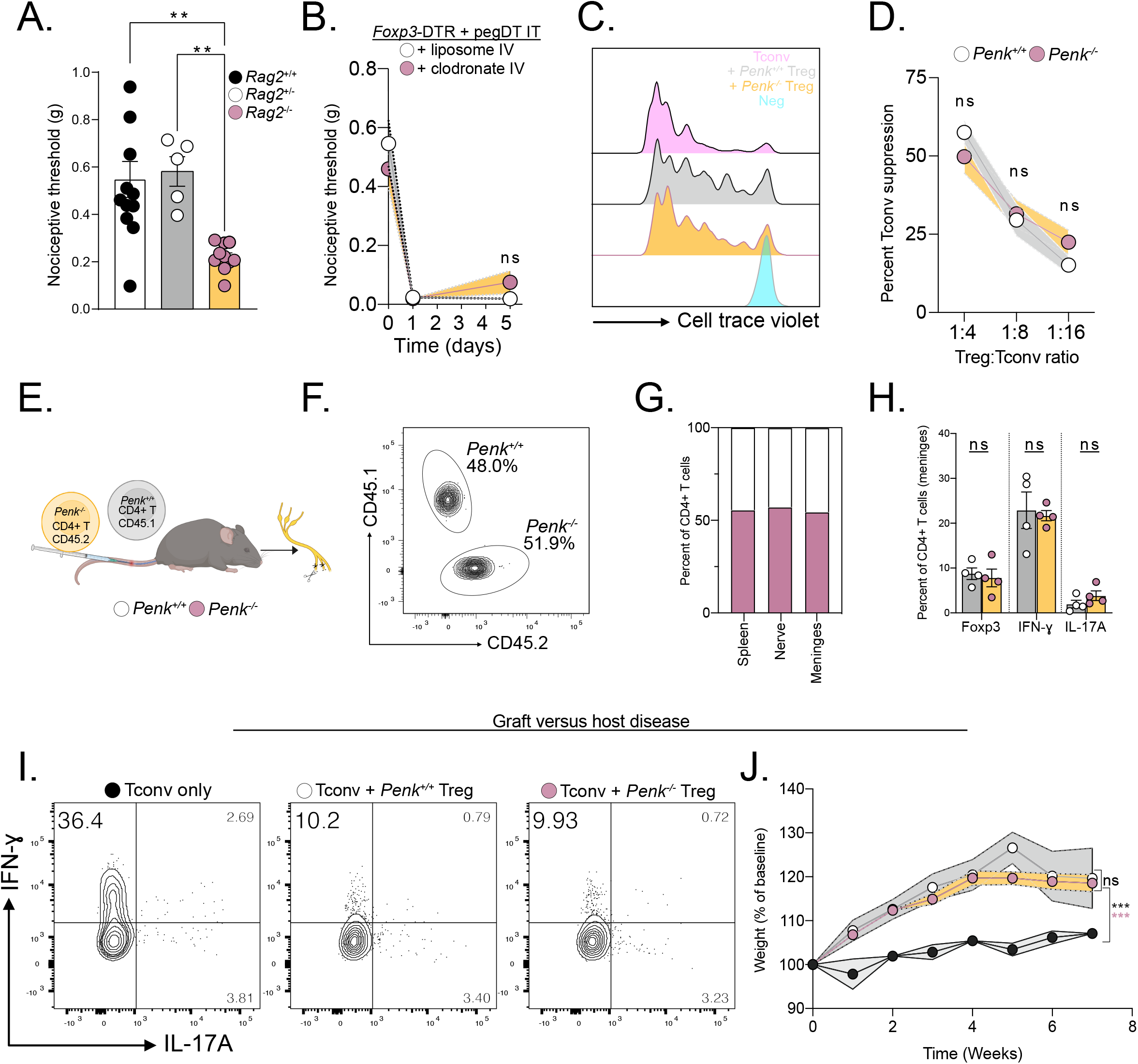
Treg-derived enkephalin is dispensable for suppressing inflammation. (A) Baseline nociceptive thresholds of uninjured *Rag2^+/+, +/-or-/-^*female mice. (B) Nociceptive thresholds of female *Foxp3*-DTR mice injected with pegDT IT + IV clodronate (pink) or control (white) liposomes showing peripheral macrophages do not mediate the nociception induced by mTreg depletion, n=5 per group. (C) Representative flow cytometry histograms of proliferated conventional T cells (Tconv) alone (pink) or 4:1 with WT Tregs (gray), *Penk^-/-^*Tregs (yellow), or unstimulated, un-proliferated cell trace violet-stained control (blue). Histogram shows cells that have not proliferated. (D) Suppression of Tconv cell proliferation by different concentrations of WT Tregs (white) or *Penk^-/-^* Tregs (yellow). (E) Schematic representation of competition experiment showing 1:1 transfer of WT or *Penk^-/-^*T cells into *Rag2^-/-^* mice. SNI surgery was performed and organs were harvested 4 weeks later for F-H. (F) Equal competition of *Penk* sufficient CD45.1 and *Penk* deficient CD45.2 CD4^+^ T cells in the meninges represented as a concatenated flow cytometry plot, n=4 per group. Representative flow cytometric plots. (G) Pooled proportion of *Penk* sufficient CD45.1 and *Penk* deficient CD45.2 CD4^+^ T cells in different organs, n=4 per genotype. (H) Percent of FoxP3^+^, IFN-^+^ and IL-17A^+^ CD4^+^ T cells from G. (I) Representative flow cytometric plots of cytokine secretion by CD4^+^ T cells after GVHD induced by transfer of pre-activated Tconv alone or combined with *Penk^+/+^*or *Penk^-/-^* Tregs. (J) Weight curves of GVHD mice, n=3-4 per group. ns = not significant, \**p*<0.05, \*\**p*<0.01,\*\*\**p*<0.001. **Related to Figure S5.**

We also used the *Penk*DTR^Δheme^ bone marrow chimeric mice to deplete all bone marrow-derived enkephalin lineage cells and assessed their contribution to the regulation of immune responses. Unlike *Foxp3-*DTR mice, we observed no changes in mouse weight or spleen size in *Penk*DTR^Δheme^ bone marrow chimeric mice chronically injected with systemic DT **(Figure S5D-E)**. Furthermore, we also did not observe any specific alterations in CD4^+^ T cell cytokine production after nerve injury in the meninges or lymphoid organs. We conclude, therefore that peripheral enkephalin does not contribute to T cell-driven inflammatory responses **(Figure S5F-G)**.

We next investigated the contribution of Treg-derived enkephalin in the regulation of conventional T cell proliferation. We assessed T cell suppression capacity by co-culturing naïve conventional CD4^+^ T cells with either WT *Penk^+/+^* or *Penk*^-/-^ Tregs. We observed no difference in the suppressive capacity of *Penk^-/-^*Treg compared to control Tregs (**Figure 5C-D**). Next, we transplanted equal amounts of CD45.1 *Penk^+/+^* or CD45.2 *Penk^-/-^* CD4^+^ T cells into *Rag2^-/-^* mice and performed SNI to measure chimerism of congenic markers among CD4^+^ T cells. We did not observe a competitive advantage or disadvantage amongst *Penk^-/-^* CD4^+^ T cells across various tissues (**Figure 5E-G**). Restimulating harvested T cells from distinct tissues with PMA/Ionomycin revealed no differences in T cell differentiation across Tregs, T helper 1 (Th1) and Th17 subsets between the *Penk-*sufficient or deficient T cells (**Figure 5H and Figure S5H-I)**. In addition, we noticed no difference in weight, health, or spleen size between *Penk*^Δheme^ *and WT*^Δheme^ bone marrow chimeric mice further revealing that peripheral enkephalin has a very limited, if any, role in the suppressing of systemic inflammatory responses **(Figure S5J-K)**.

Finally, using an adoptive transfer-based graft versus host disease (GVHD) model, we assessed whether Treg-derived enkephalin is required for suppressing immune responses. As expected, we observed a profound Th1 response in mice transferred with activated Tconv alone. However, mice that received additional transfers of either *Penk^+/+^*or *Penk^-/-^* Tregs equally suppressed Th1 responses, reduced GVHD severity and mitigated weight dysregulation (**Figure 5 I-J and S5L-N)**. In summary, we conclude that Treg-derived enkephalin does not contribute to any inflammatory response restraint mechanism. Rather, we conclude that Tregs can suppress pain sensitivity through a mechanism that is independent of their function in immunosuppression.

### Delta opioid receptor signaling on MrgprD^+^ primary afferent DRG neurons is required for the anti-allodynic function of mTregs

Enkephalin is a potent agonist at the δOR, and, to a lesser extent the μOR. In our previous studies, we demonstrated the divergence of expression and function of δOR and μOR in mediating distinct pain modalities. Specifically, δOR is expressed on nonpeptidergic IB4^+^ unmyelinated as well as myelinated primary afferents and selectively regulates mechanical thresholds and nerve injury-induced mechanical hypersensitivity^21^. Conversely, the μOR is expressed on Trpv1^+^ nociceptors and selectively regulates thermal hyperalgesia. In addition, a spinal δOR can dampen mechanical hypersensitivity by inhibiting the excitability of somatostatin-positive dorsal horn interneurons^23^.

To assess the requirement of PNS or CNS δOR circuits in coordinating the anti-allodynic effect of mTregs, we intravenously injected *Oprd1^+/+^* control or *Oprd1^fl/fl^* mice with AAV.PHP.S-CAG-Cre or AAV.PHP.eB-CAG-Cre. This approach selectively introduces Cre recombinase and targets deletion of δOR into to the PNS (DRG) or CNS (spinal cord and brain), respectively (**Figure 6A**). Three weeks after the AAV injection, we performed SNI and four weeks later administered IL-2. Mice selectively lacking δOR in the PNS lost the capacity to respond to the anti-allodynic effect of IL-2, but the effects of IL-2 were preserved in mice lacking δOR in the CNS (**Figure 6B-C**). We conclude that a sensory neuron-expressed, presynaptic δOR coordinates mTreg suppression of mechanical pain hypersensitivity.

**Figure 6.**
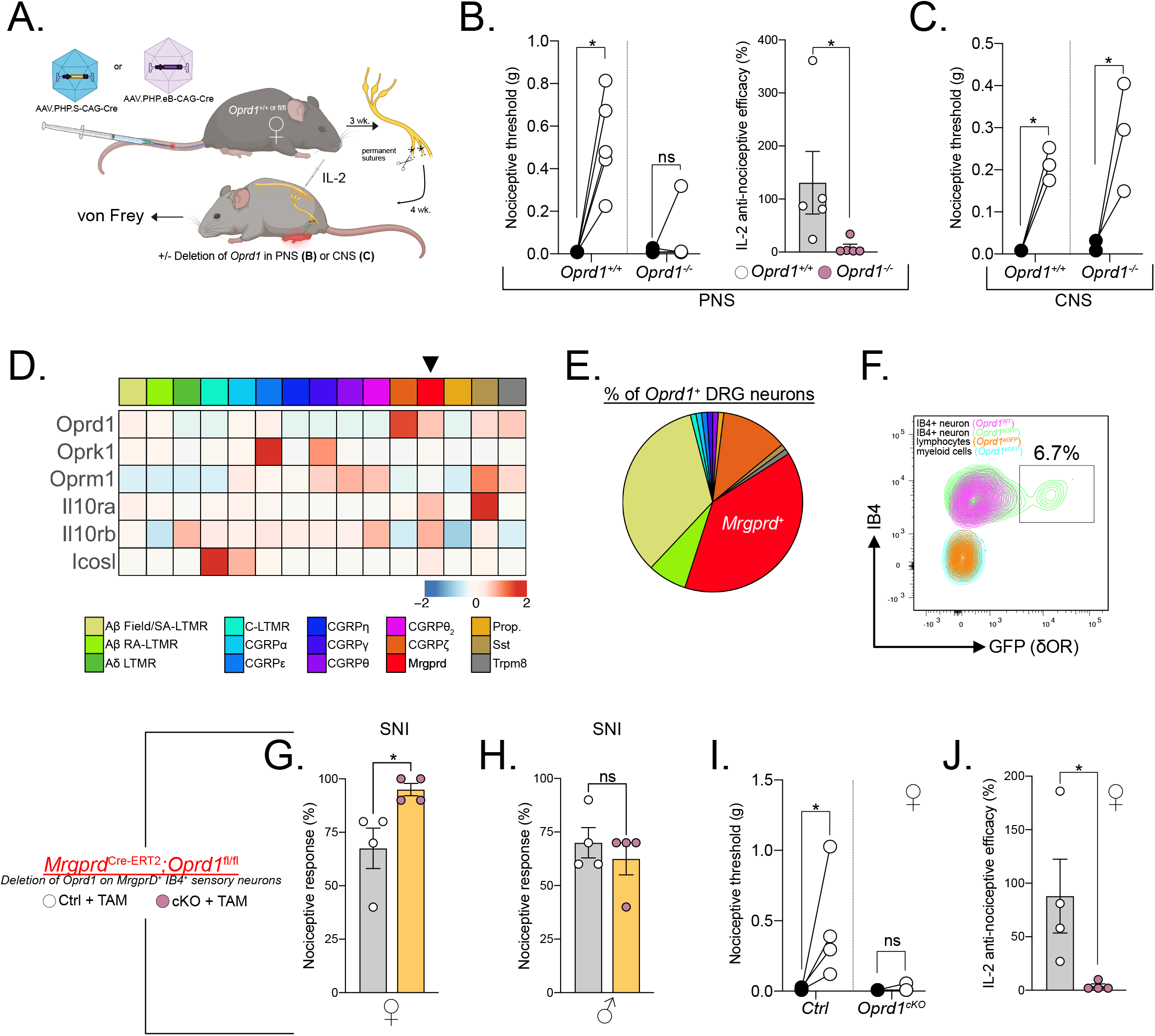
δOR on MrgprD^+^ sensory neurons is required for mTreg mediated anti- nociception. (A) Schematic representation of AAV-induced ablation of *Oprd1* in the PNS (B-C) or CNS. (B) Nociceptive thresholds of female mice lacking *Oprd1* in the PNS after mTreg expansion compared to controls. (C) Anti-nociceptive efficacy of mTreg expansion. (D) No difference in nociceptive thresholds in female mice lacking *Oprd1* in the CNS after mTreg expansion compared to controls. (E) Heatmap of row normalized expression from DRG sensory neurons clusters from combined GSE139088 and GSE201653. (F) Proportions of sensory neuron clusters expressing *Oprd1* from E. (G) Representative flow cytometry plot of δOR-GFP (green) expression on IB4^+^ MrgprD^+^ DRG sensory neurons compared to cells from non-reporter mice (purple). Overlaid are lymphoid CD45^+^ CD90.2^+^ cells and myeloid CD45^+^ CD11b^+^ cells from the DRG. (H-I) Percent response to 0.008 g von Frey fiber stimulation after SNI in female (H) or male (I) mice conditionally lacking δOR on MrgprD^+^ neurons (pink) or controls (white). (J) Nociceptive thresholds after mTreg expansion in female mice lacking *Oprd1* in MrgprD^+^ neurons compared to controls. (K) Anti-nociceptive efficacy of mTreg expansion. SA-LTMR= Slowly adapting low-threshold mechanoreceptor, RA-LTMR= rapidly-adapting low-threshold mechanoreceptor, MrgprD= Mas-related G protein-coupled receptor D, Prop.= proprioceptor, SST: somatostatin, Trpm8= Transient receptor potential cation channel subfamily M member 8, TAM= Tamoxifen, cKO= Conditional KO, ns = not significant, \**p*<0.05, \*\**p*<0.01,\*\*\**p*<0.001.

Next, we identified the specific sensory neuron subset that coordinates the anti- nociception mediated by mTreg-derived enkephalin. Previous studies of δOR expression on DRG sensory neurons using *Oprd1^eGFP^*reporter mice revealed that approximately half of the reporter-positive cells in the DRG are myelinated neurons, while approximately 36% are IB4^+^ non-peptidergic neurons expressing the MrgprD receptor. Using established single cell RNA sequencing resources, we found that the MrgprD^+^ subset of DRG sensory neurons not only expresses *Oprd1*, but also other receptors for Treg ligands, including *Il10ra and Icosl*, which have been implicated in suppression of pain thresholds^46–48^ (**Figure 6D**). The total proportion of sensory neurons expressing the *Oprd1* transcript matches previous data using *Oprd1^eGFP^*reporter mice (**Figure 6E**). Our subsequent flow cytometry-based profiling of *Oprd1^eGFP^* reporter expression on DRG cells confirmed GFP expression specifically on IB4^+^ CD45^-^ Thy1^+^ sensory neurons, which corresponds to the MrgprD^+^ nonpeptidergic nociceptive neuron population. Importantly, the flow analysis confirmed the absence of GFP expression on broadly defined CD45^+^ CD90.2^+^ IB4^-^ lymphoid cells and CD45^+^ CD11b^+^ CD90.2^-^ IB4^-^ myeloid cells (**Figure 6F**). We also confirmed the absence of GFP expression on microglia as well as on immune cells profiled from the draining lymph nodes (data not shown)^23^. Based on this selective Oprd1 expression profile, we generated mice in which MrgprD^+^ neurons lack the δOR (*MrgprD*^Cre-ERT2^;*Oprd1*^fl/fl^). Female *MrgprD*^Cre-ERT2^*;Oprd1*^fl/fl^ mice, but not their male counterparts, exhibited exaggerated mechanical hypersensitivity after SNI compared to tamoxifen-injected sex-matched littermate controls (**Figure 6G-H**). Female mice lacking δOR on MrgprD^+^ sensory neurons and treated with IL-2 IT four weeks following SNI displayed a complete deficiency in IL-2 anti-allodynic efficacy (**Figure 6I-J**). We conclude that, the enkephalin receptor δOR, expressed specifically by MrgprD^+^ sensory neurons, mediates the anti-nociceptive function of mTregs.

## Discussion

In this report, we describe a novel, sexually dimorphic mechanism for pain regulation by the immune system. Using a range of site-selective targeting strategies to deplete or expand Tregs within the recently recognized borders of the nervous system, specifically the meninges, we find that meningeal Tregs (mTregs) can profoundly modulate mechanical hypersensitivity. Strikingly, this pain regulatory function of mTregs is sex-specific and controlled by gonadal hormones. Although proenkephalin expression by Tregs has been observed in sequencing studies, its functional relevance to nociception had not been explored. Here, we demonstrate that enkephalin secreted by mTregs acts on δ-opioid receptors (δOR) on primary sensory neurons to selectively modulate mechanical sensitivity. Our findings provide the first mechanism of Treg-mediated suppression of nociception and establish regulatory T cells as key sentinels of pain homeostasis.

To assess for sex differences in transcriptional identify after nerve injury, primary afferent neurons, including those expressing MrgprD, have recently been sorted for deep RNA sequencing^52^. Surprisingly, very few differences were found between the sexes, suggesting a lack of a strong intersection of sex and injury in the transcriptional identity of peripheral neurons. The lack of transcriptional differences may also suggest that non-neuronal cells may be a primary determinant of sex-selective pain modulation. Indeed, preclinical research has demonstrated the involvement of T cells in driving pain phenotypes in female mice^53^. Additionally, human leukocyte antigen (HLA) risk alleles have been identified for human chronic pain conditions, further suggesting a potential role for T cells in pain modulation^24,54^.

Regulatory T cells display broad tissue supportive roles that extend beyond their originally described function in suppressing inflammation^8,11,12,20^. A major advantage of our analysis is that we utilized a site-selective mTreg ablation strategy that preserved peripheral Tregs and avoided systemic inflammation. Using this approach, we identify a novel, sex-specific mechanism by which Tregs modulate nociceptor activity to regulate pain sensitivity in the context of health and nerve injury. Although, proenkephalin- expressing Tregs have been identified in various tissues, their functional assessment has been limited^19,37,38^. Somewhat paradoxically, a recent pre-print study demonstrated a small but statistically significant decrease in basal heat sensitivity in both female and male mice conditionally depleted for *Penk* expression in systemic Tregs. Mechanical thresholds and other sensory modalities were however not examined^55^. In distinct contrast, we uncover a sensory modality selective function of mTreg that is consistent with prior findings of δOR agonism, namely providing relief of mechanical but not heat pain^21,56^. Whether mTregs tonically restrain nociception is difficult to conclude. Our finding of increased basal sensitivity in *Rag2^-/-^* or *Penk*^Δheme^ mice, is supportive of this hypothesis, however, alternative possibilities exist. It is conceivable that mTreg deficiency could lead to inflammation, which may alter nociceptor sensitivity. Additionally, the endogenous opioid signaling pathway may have a role in opioid induced analgesia. It is significant that naloxone, a non-selective opioid receptor antagonist IV injection, does not induce pain in healthy individuals. Whether experiments in uninjured female mice have been performed is unclear^57^.

Our observations suggest that sex hormones, rather than sex chromosomes, are the main drivers of mTreg-induced anti-nociception. While the specific sex hormones involved in regulating mTreg function in pain remain unclear, previous studies have implicated estrogen and progesterone in modulating neuropathic pain associated with SNI^49^. Estrogen administration increases *Penk* expression in the whole spinal cord^58^. Additionally, in CD4^+^ T cells, the estrogen receptor engages conserved non-coding sequences (CNS) in *Foxp3* enhancer regions^59^. Importantly, we reveal an anti- nociceptive role for regulatory T cells that is distinct from their well-established functions in immune suppression and tissue repair. Furthermore, we demonstrate that this mechanism operates within nervous system tissues, at a site distant from the peripheral nerve injury, highlighting the immune system’s remarkable ability to modulate nociception.

## Methods

### Mice

All mouse experiments were approved by UCSF Institutional Animal Care and Use Committee and conducted in accordance with the guidelines established by the Institutional Animal Care and Use Committee and Laboratory Animal Resource Center. All mice experiments were performed on age-matched adult male and female mice at a starting age between 8 and 14 weeks old. Littermate controls were used for all experiments when feasible. Mice were bred in-house and backcrossed over 10 generations to C56BL/6 breeders obtained from Jackson labs. Experimental mice were co-housed to maintain the same microbiome. They were maintained in a temperature (21°C) - and light (12h light/dark cycle)-controlled environment and were provided with food and water *ad libidum*. The following mouse strains are used: C57BL/6J (JAX #000664), *Foxp3*-DTR (B6.129(Cg)-Foxp3tm3 (Hbegf/GFP)Ayr/J, JAX# 016958), *Foxp3*^eGFP-Cre-ERT2^ (Foxp3tm9 (EGFP/cre/ERT2)Ayr/J; JAX#016961), Ai9 (B6.Cg- *Gt(ROSA)26Sor^tm14(CAG-tdTomato)Hze^*/J; JAX#007914), Four Core Genotypes (B6.Cg-Tg(Sry)2Ei Srydl1Rlb T(XTmsb4x-Hccs;Y)1Dto/ArnoJ; JAX#010905), Penk-IRES2- Cre(B6;129S-Penktm2(cre)Hze/J; JAX#025112), Rosa26-lSL-iDTR (CBy.B6- Gt(ROSA)26Sortm1(HBEGF)Awai/J;JAX#008040),CD45.1 (B6.SJL-Ptprca Pepcb/BoyJ; JAX# 002014), B6C3F1/J (JAX#100010), *Rag2*^-/-^ (B6.Cg-Rag2tm1.1Cgn/J; JAX#008449), Mrgprd-CreERT2 (Mrgprdtm1.1(cre/ERT2)Wql/J; JAX# 031286), Oprd1fl/fl (B6;129-Oprd1tm1.1Cgrf/KffJ; JAX# 030075), DOR-eGFP (B6;129S2-Oprd1tm2Kff/J; JAX#029012). *Penk^-/-^* were kindly provided by Dr. John Pintar on a C57BL/6 background (MGI 3628668)^60^. *Foxp3*-DTR (X-linked) was mated with male Four Core Genotypes XY^Sry-^Sry^Tg^ and the X^Foxp3-DTRYSry-^Sry^Tg^ male mice were mated to homozygous *Foxp3*-DTR female mice.

### Bone marrow transplantation

CD45 mismatched host recipient mice were irradiated at 550 cGy twice, 5 hours apart and injected retro-orbitally with 5x10^6^ cells from the bone marrow of CD45.1 WT, *Penk*^Cre^;*Rosa26*^DTR^ *Penk^-/-^* or a 1:1 mix of *Foxp3*-DTR and *Penk^-/-^* sex-matched 7-10 week old mice. Mice were kept on doxycycline chow for the first week (Bioserv #S3888) and chimerism was assessed at 8 weeks post-transplant.

### Pharmacological interventions

Animals were randomly assigned to vehicle control or treatment group. Pegylated diphtheria toxin (pegDT) was a generous gift from Ana I. Domingos and generated as previously described^61^. pegDT (20 ng) or corresponding phosphate-buffered saline (PBS) control were injected intrathecally (IT) in a volume of 5µL in naive mice below lumbar level L4. All intrathecal injections were performed in non-anesthetized, lightly restrained mice and injections were validated by a sudden flick of the tail. Of note, the 5- 10 µL injection distributes predominantly to the lumbo-sacral cord given lidocaine IT injection at that segment paralyzes the hindpaws but not forepaws. Non-pegylated diphtheria toxin (30 ng/g, Sigma Cat#322326) or corresponding saline control were administered in a volume of 200 µl every three days intraperitoneally (IP). IL-2 (0.1 µg, Peprotech, Cat# 212-12) or PBS vehicle control were administered daily for three consecutive days IT. Selective δOR agonist, [D-Ala2]-Deltorphin 2 (15 µg, Abcam Cat #ab120708, CAS 122752-16-3) and selective δOR antagonist, naltrindole (5 µg, Sigma Cat# N115) administrations were performed 30 min before behavior experiments.

### Tamoxifen injections

*Mrgprd*^CreERT2^;*Oprd1*^fl/fl^ and *Foxp3^eGFP-Cre-ERT2^;Rosa26^tdTomato^*mice were injected IP with tamoxifen (Sigma Cat #5648) 100 mg/kg in corn oil (Sigma Cat #8267) for five consecutive days to induce Cre-mediated recombination.

### Intrathecal dye tracing

Naive mice were intrathecally injected with 5µL of Evans Blue 1% solution (Sigma, Cat #E2129) or pegylated DyLight 650-4xPEG NHS Ester (Thermo Fischer, Cat #62274). 24 hours post-injection, mice were anesthetized with avertin and euthanized by decapitation. Spinal cord and brain meninges, spinal cord, brain, dorsal root ganglia and trigeminal ganglia, sciatic nerve and lymph nodes were assessed for dye uptake.

### Animal behavior

For all behavioral tests, the experimenter was blind to genotype and treatment and performed during the light cycle. The project utilized both male and female experimenters but a predominant number of experiments were performed by a female investigator^62^.

### von Frey measurement of mechanical hypersensitivity

Mice were acclimatized once to the von Frey apparatus for two hours. The lateral plantar surface of the ipsilateral and contralateral hind paws (sural innervation) was stimulated with von Frey hairs of logarithmically increasing stiffness (Stoelting Cat # 58011). Animals were habituated on a wire mesh for 1 hour, after which they were tested with von Frey filaments (0.008, 0.02, 0.04, 0.07, 0.16, 0.4, 0.6, 1, 1.4, 2, 4 and 8 g) using the Dixon up–down method^63,64^. The von Frey hairs were held for 3 sec with intervals of several minutes between each stimulation. For the Dixon up-down method we recorded 2 days of baseline mechanical sensitivity which were averaged. After SNI, nearly all mice reached a 50% paw withdrawal threshold of the lowest filament (0.008 g), thus we utilized a single fiber method of testing to achieve resolution of allodynia severity. Mice were stimulated 10 times with the 0.008 g filament. The filament was applied for 3 seconds and the number of positive responses across the 10 stimulations were registered as percent nociceptive responses.

### Hargreaves measurement of heat hypersensitivity

Mice were acclimatized for 30 min in plexiglass cylinders. The mice were then placed on the glass of a Hargreaves apparatus and the latency to withdraw the paw from the heat source was recorded. Each paw was tested three times and latencies were averaged over the trials.

### Acetone induced cold sensitivity

Mice were habituated for 60 min on a mesh in plexiglass cylinders. A syringe was used to spray 50 µl of acetone (Thermo Scientific Cat # 423240010) onto the plantar surface of the paw and the behaviors were video recorded for 30 seconds after each trial using a Sony HDR-CX440 camera. The left hind paw was tested five times and positive responses included withdrawals, shakes, licks and jumps. Results are displayed as the total number of behaviors across the five trials.

### Tail flick measurement of heat hypersensitivity

Mice were placed in a restrainer and 2 cm of the tip of the tail was submerged in a 52°C water bath. The latency (seconds) to withdraw the tail from the water was recorded. A cut-off of 15 s was set to prevent tissue damage and testing was performed with intervals of several minutes between each stimulation. Mice were tested three times and withdrawal latencies were averaged.

### Hot plate measurement of heat hypersensitivity

Mice were acclimated to the testing environment as described above. The hot plate temperature was set to 52°C. The mouse was placed on the plate and the latency to shake, lick or bite a hindpaw was scored. A cut-off of 20 s was set to prevent tissue damage.

### Pin prick withdrawal test

Mice were habituated for 60 min on a mesh in plexiglass cylinders. A 27G needle was gentle applied onto the hindpaws, with minutes between each stimulation for a total of 5 stimulations per paw. Mice were scored as follow: 1: brief withdrawal, 2: mice lifting their paw for 1 sec, 3: mice lifting their paw for 2 sec or shake, 4: lick. Results are displayed as the total number of behaviors across the five trials.

### Brush withdrawal test

Mice were habituated for 60 min on a mesh in plexiglass cylinders. A 5-0 brush was gently applied onto the hindpaws, with minutes between each stimulation for a total of 5 stimulations per paw. Mice were scored as follow: 1: brief withdrawal, 2: mice lifting their paw for 1 sec, 3: mice lifting their paw for 2 sec or shake, 4: lick. Results are displayed as the total number of behaviors across the five trials.

### Rotarod

Mice were acclimatized to the testing room and trained by placing them on an accelerating rotarod for a maximum of 60 seconds at low speed, three times with training taking place on two consecutive days. Latency to fall was measured with a cutoff of 300 seconds. The procedure was repeated three times and latencies averaged across trials.

### Spared Nerve injury

We employed an established, robust and reliable spared nerve injury (SNI) model to induce a chronic neuropathic injury^65^. This model utilizes non-healing surgical intervention on two branches of the sciatic nerve (the common peroneal and tibial branches), while sparing the third branch (the sural branch) for sensory testing on the lateral portion of the hindpaw. Briefly, mice were anesthetized with isoflurane (3% for induction and 1.5% for maintenance, mixed with oxygen). The fur on the left hind leg was shaved and disinfected with 3 passages of alcohol and iodine solution, alternatively. A 1 cm incision was performed on the upper thigh skin, near the division point of the sciatic nerve. A 2% lidocaine solution was applied and the biceps femoris muscle was gently separated through a blunt opening to reveal the sciatic nerve’s common peroneal, tibial, and sural branches. The common peroneal and tibial nerves were ligated with non-dissolvable 8-0 silk sutures (Fine Science Tools Cat # 12052-08). Subsequently, a 2 mm segment from both the common peroneal and tibial nerves was transected, ensuring the sural nerve remained undisturbed. The muscle and the skin were stitched using 6–0 sutures (Henry Schein Surgical suture Cat #101-2636), and the skin was further sealed with a tissue adhesive (3M Vetbond Cat # 1469SB), after an ethanol solution application. Mice were kept on heating pad until they regained consciousness and demonstrated stable, balanced locomotion. Mice were transferred into their home cage and observed meticulously for the next two days.

### Immunohistochemistry

Avertin-anesthetized mice were transcardially perfused with 10 ml of 1× PBS followed by 30 ml of 4% paraformaldehyde (PFA, Thermo Scientific Cat # 119690010) diluted in PBS. After perfusion, spinal cord, sciatic nerves, lymph nodes, spleens, brains and DRG were collected, postfixed in 4% PFA solution at 4°C for 5 h and then cryoprotected in 30% sucrose in PBS at 4°C.

Spinal meninges were harvested from fixed spinal cords. Spinal cords were transferred in PBS and meninges were gently peeled into a single sheet onto a microscope slide after a longitudinal hemisection of the spinal cord. Brain meninges were similarly harvested from the skull. Frozen tissues were embedded at −35°C in O.C.T. compound and 30 µm transverse spinal cord sections were generated using a Leica SM220R sliding microtome and 20 µm DRG sections were generated using a cryostat (Thermo Fisher Scientific) on SuperFrost Plus slides. Spinal cord sections were processed as free-floating. Sections were blocked (10% NGS, 1% BSA, 0.05% Tween-20, 0.1% Triton X-100 in PBS) and incubated in 0.3 M glycine containing 0.2% Tween 20. Sections were labeled in blocking buffer for 24 hours at 4°C. Slides were coverslipped with Fluoromount-G (Thermo Fisher Scientific). Fluorescence images were acquired using an Olympus FV3000 confocal microscope and quantified using ImageJ (Fiji).

### Tissue clearing

Whole DRG or spinal cords from *Foxp3*^eGFP-Cre-ERT2^;Ai9 mice were cleared after PFA fixation using SHIELD tissue clearing (LifeCanvas PCK-500)^66^. Tissues were washed in PBS then processed as previously described^66^. Briefly, the tissues were incubated in epoxy solution (SHIELD OFF) for 10 hours at 4°C with gentle shaking then incubated overnight at 37°C in SHIELD ON-Epoxy solution for epoxy crosslinking. DRG were then further incubated in SHIELD ON solution for 10h and delipidated for two days (DRG) to five days (spinal cord) at 45°C with shaking then washed with PBS. Whole mount DRG and spinal cords were acquired in FocusClear reflexive index matching solution (CelExplorer, FC-102).

### Tissue digestion

Mice were injected intravenously with 50 µl of anti-ARTC2 nanobody (Biolegend Cat # 149802) in 200 µL of PBS 30 minutes before euthanasia to protect Treg during harvest from purinergic-mediated cell death^67^. 5 minutes before harvest mice were injected intravenously with 6 µg of FITC-conjugated anti-CD45 antibody to label blood immune cells in 200 µL of PBS ^27^. Avertin-anesthetized mice were decapitated, and spinal cord meninges, brain meninges, L4-6 DRG, sciatic nerves, lymph nodes, brains and spleens were harvested. Spinal cord meninges, brain meninges, DRG, sciatic nerves and brains were crushed with the back of a 3ml syringe in a serrated 24 well plate and triturated in digestion media (Liberase TM (0.208 WU/ml) (Roche Cat # 054010200001) and DNase I (40 ug/ml) (Sigma Cat # DN25) in 1.0 ml cRPMI (RPMI supplemented with 10% (vol/vol) fetal bovine serum (FBS), 1% (vol/vol) HEPES, 1% (vol/vol) Sodium Pyruvate, 1% (vol/vol) penicillin-streptomycin). They were digested for 30 min at 37°C, 220 RPM and triturated every 15 minutes. Digested samples were again triturated and passed over a 40 µm cell strainer and any remaining tissue pieces mashed through the cell strainer. Cell strainers were flushed with staining media (PBS w/o Mg^2+^ and Ca^2+^ supplemented with 3% FBS, 2 mM EDTA and 0.05% NaN_3_). Single-cell suspensions were centrifuged at 500 g at 4°C, washed and resuspended in staining media. Spleens and lymph node immune cells were obtained by mashing the tissues over a 40 µm cell strainer and washed with staining media.

### Cell stimulation

Isolated single cell suspensions were incubated for 4 hr at 37°C in complete IMDM (supplemented with 10% FBS, 2 mM L-glutamine, 10 mM HEPES, 1% sodium pyruvate, 1% penicillin-streptomycin, 50 µM 2-mercaptoethanol) with (phorbol 12-myristate 13- acetate) PMA, Ionomycin in the presence of Brefeldin A and Monensin (Tonbo, TNB- 4975).

### Flow cytometry

Single-cell suspensions were stained in a 96 well plate. Briefly, they were washed with 250 µL of staining media and stained with viability dye (1:500) and cell surface antibodies (1:100) in 100 µL of staining media with Fc shield (1:100). Samples were washed twice in staining media and stained for intracellular cytokines or for Foxp3 according to manufacturer recommendations with BD Cytofix/Cytoperm Fixation/Permeabilization Kit (BD 554714, AB_2869008). Samples were further stained with conjugated intracellular antibodies (1:100) overnight at 4°C in BD permeabilization/wash buffer. Samples were washed with permeabilization/wash buffer twice and resuspended in 200 µL of staining media. For visualization of both tdTomato reporter signal and intranuclear Foxp3 signal, cells were fixed in 200 µL of freshly prepared 2% formaldehyde (EM grade) in PBS for 60 minutes exactly and washed with eBioscience Permeabilization buffer (eBioscience Foxp3 perm-kit) and stained overnight in 1x eBioscience Permeabilization buffer at 4°C^68^. Cells were washed twice in the buffer and resuspended in staining media. For Met-enkephalin staining, we screened multiple commercially available antibodies and selected an antibody that showed positive staining in wildtype mTreg but not *Penk^-/-^* mTreg. The antibody was conjugated to fluorescent phycoerythrin with Lightning-link conjugation kit (ab102918) and utilized after cell stimulation and intracellular cytokine staining. Cells were counted with 50 µL of counting beads (Thermo Fisher Scientific Cat # C36950) and samples were analyzed using a BD FACSCanto2 or BD FACS Aria Fusion flow cytometer (BD Biosciences). Positive and negative selection gates were set using fluorescence minus unstained cells. For negative control of enkephalin staining, *Penk^-/-^* samples were used for gating. Fluorescence intensity distribution was analyzed using FlowJo 10 software (BD Biosciences). Antibodies for flow cytometry are listed in the resource table. Lineage exclusion markers include viability dye, CD11b (to exclude myeloid cells), B220 (to exclude B cells), Ter119 (to exclude red blood cells).

### Cell sorting

Spleens were mashed on a cell strainer and cells were pelleted at 500 g for 10 min at 4°C. Cells were stained for viability and lineage exclusion markers, in addition to CD45, CD4, CD45RB to stain naïve Tconv and CD25 to stain Tregs. Cells were pelleted, washed and incubated with CD4 negative selection beads and purified on a LS magnetic column (Miltenyi Biotec). Cells were double sorted in BD FACS Aria Fusion for Singlet, Live, CD45^+^ CD4^+^ CD45RB^+^ CD25^-^ for Tconv or Singlet, Live, CD45^+^ CD4^+^ CD25^+^ CD45RB^-^ for Treg into complete IMDM.

### T cell suppression assay

Sorted Tconv from lymphoid organs of CD45.1 female mice were labeled with cell trace violet (Thermo-Fisher, Scientific #C34571) according to manufacturer instructions.

0.25x10^5^ Tconv per well were cultured with distinct dilution of Treg from CD45.2 WT or *Penk* deficient lymphoid organs. Cells were then washed and resuspended with mouse anti-CD3/CD28 Dynabeads at a 1:1 ratio of beads to Tconv. Cells were incubated for 96 hours in a humidified incubator at 37°C. Cells were washed and resuspended in staining media and suppression ratio was calculated by dividing the percent proliferated cells from incubated Tconv^+^ Treg samples by percent proliferated cells from Tconv only samples^69^.

### T cell adoptive transfer

#### Competition assay

1x10^6^ negatively selected bulk CD4^+^ T cells from WT female and *Penk* deficient lymphoid organs were transplanted into female *Rag2^-/-^* mice. SNI was performed and organs were collected 28 days later for cell stimulation and flow cytometry.

#### Graft versus host disease (GVHD)

GVHD was established as previously described. Briefly, WT or *Penk* deficient female mice bone marrow was transplanted into MHC mismatched B6C3F1/J female mice to activate T cells. 0.25x10^6^ sorted WT Tconv were transplanted into male *Rag2^-/-^* mice to induce chronic GVHD either without Tregs or in the presence of 0.125 x10^6^ WT or *Penk* deficient Tregs. Mice were measured for weight changes and GVHD score. Mice were euthanized then harvested for cytokine secretion assay by flow cytometry^70,71^. GVHD scoring is as follows: 0 = no signs of GVHD, 1 = visible signs of GVHD (hunching, lethargy, ruffled fur), 2 = no weight gain, 3 = 0-5% weight loss, 4 = >5% weight loss. One Tconv mouse did not survive for harvesting for cytokine stimulation.

### Analysis of sequencing data

#### Bulk RNA-seq

Raw files GSM4677053-064 from GEO dataset GSE154680, and all files from GSE GSE180020 were gathered and aligned using STAR for uniquely mapped reads (outFilterMultimapNmax 1–outFilterMatchNmin 30–alignIntronMin 20–alignIntronMax 1- 0000). Data was annotated with GENCODE GRCm38/mm10 genome assembly. Raw count tables were normalized by median of ratios method with DESeq2 package from Bioconductor to analyze for differential expression.

#### ATAC-seq

Fastq files were gathered from SRR12264679-94 from GSE154680. Raw reads were mapped to the mouse mm10 genome assembly using STAR alignment (-- outFilterMismatchNoverLmax 0.04 --outFilterMismatchNmax 999 -- alignSJDBoverhangMin 1 --outFilterMultimapNmax 1 --alignIntronMin 20 -- alignIntronMax 1000000 --alignMatesGapMax 1000000). Bam files were generated by STAR. PCR duplicates were removed by Picard, and peak calling performed using MACS2 (--keep-dup 1 --bw 500 -n output --nomodel --extsize 400 --slocal 5000 --llocal 100000 -q 0.01) PMID: 22936215). To generate bigwig files for ATAC-seq datasets, all aligned bam files were merged by condition using samtools merge. Bedtools genomecov was run to covert the merged bam files into a bedgraph files. Finally, bedGraphToBigWig (ucsc-tools/363) was used to generate the bigwig files displayed on browser tracks using the IGV browser and compared to existing encode ATAC and Chip-Seq peaks.

#### scRNA-seq

Fastq files were gathered from GEO from datasets GSE139088 GSE201653 and initial counts were obtained using the Cell Ranger pipeline^48,72^. Using Seurat v4, individual cells were removed from the data set if they had fewer than 1000 discovered genes/features, fewer than 1000 UMI or greater than 10% reads mapping to mitochondrial genes. 2000 variable genes were found for each normalized library, and anchors were selected for integration with dimensionality of each dataset set at 30. Glial cells noted for the markers of *Sparc* and *Mpz* and non-neuronal cells lacking the expression of Avil were excluded. Variable genes were identified from the merged dataset, and PCA and UMAP were ran to generate new UMAP coordinates with a dimensionality of 30 and clustering was performed with a resolution of 0.5.

Findallmarkers function utilizing a Wilcoxon rank-sum test was used to find cluster specific markers and annotation was performed as recently established^48^. Number of *Oprd1* expressing cells were defined by a threshold of non-zero expression.

## Statistical analysis

Statistical analysis was performed using GraphPad Prism 9 software. Data are presented as mean ± SEM. Differences pre- and post-injection within a single group were assessed using a Wilcoxon matched-pairs signed rank test. Differences between two groups were assessed using a Mann-Whitney test. Statistical analysis for multiple comparisons were performed using Kruskal-Wallis test followed by Dunn’s multiple comparison test or a Two-Way ANOVA followed by Sidak’s multiple comparison test. *p* < 0.05 (*), *p* < 0.01 (**), *p* < 0.001 (***).

## Acknowledgement

We thank Dr. Dena Dubal and her laboratory for the FCG mouse, Dr. Kevin Yackle for *Penk*^Cre^ mice, Dr. Ari Molofsky for *Foxp3*^Cre-ERT2^*Rosa26*^TdTomato^ mice, Dr. Amynah Pradhan for *Oprd1^eGFP^*mice and Dr. Mike Ansonoff for assistance with Penk^-/-^ tissue isolation. We thank additional members of the Basbaum laboratory and UCSF ImmunoX for critical feedback. Funding for this work was supported by grants Canadian Institute of Health Research (CIHR) (to É.M.), the Fonds de Recherche en Santé-Québec (to É.M.), the Dermatology Foundation (Career Development Award to S.W.K.), the Sandler Foundation PBBR (to S.W.K), Grunfeld Scholar Award from SFVAMC (to S.W.K), T32AR007175-44 (to S.W.K), NIH NSR35NS097306 (to A.I.B.) and Open Philanthropy (to A.I.B.). Figures were generated with BioRender.com.

## Author contributions

É.M. and S.W.K. designed experiments. É.M., B.C.M, J.B., S.R. N.P.K. W.L.E. and S.W.K. performed experiments, data analysis or visualization.

J.E.P. and A.I.D. provided critical reagents or tools. S.W.K, É.M. and A.I.B., acquired funding and provided supervision. É.M, A.I.B and S.W.K. wrote the manuscript.

## Material availability and requests

No new reagents, original code or original genomic datasets were generated. Requests for reagents or mice can be sent to.

## Diversity, equity, and inclusion statement

Authors support diversity and inclusion values. At least one author, including the lead author, self-identifies as a woman. At least one author identifies as an under-represented minority and/or as an immigrant.

## Supplemental Figure Legends

**Figure S1. Flow cytometric gating strategy of meningeal Tregs.**

(A) Gating strategy to quantify tissue Treg numbers. Numbers indicate gate frequency. Mice were injected IT with ARTC2 nanobody to minimize Treg apoptosis and injected IV with anti-CD45 FITC (pink) antibody or vehicle injected (blue) to label vascular immune cells.

**Figure S2. pegDT IT injection avoids systemic inflammation and weight loss in *Foxp3*-DTR mice.**

(A) Evan’s blue staining after IT injection showing diffusion into the cerebellum, the olfactory bulb, the cervical and lumbar lymph nodes, the spinal cord meninges and the lumbar DRG. (B) pegDyLight650 IT injection exhibits a more limited diffusion than Evan’s blue. (C) Weight curves of *Foxp3*-DTR mice injected with IT pegDT, IP DT or IT vehicle every 3 days demonstrating a lack of weight loss after site-selective Treg ablation. Arrows represent DT injections. (D and E) Representative images of spleen sizes and spleen weights from mice in (C). (F) Survival curves of mice in (C). ns = not significant, \**p*<0.05, \*\**p*<0.01,\*\*\**p*<0.001.

**Figure S3. mTreg depletion selectively induces mechanical hypersensitivity in female mice.**

(A) von Frey, (B) Hargreaves, (C) hotplate, (D) tail flick, (E) acetone, (F) pinprick, (G) brush and (G) rotarod behavioral testing in *FoxP3*-DTR mice injected with a single dose of IT pegDT. n = 4-20 mice per group. ns = not significant,\*\*\*\**p*<0.0001.

**Figure S4. mTreg expansion selectively improves mechanical hypersensitivity in injured female mice.**

(A) mTreg expansion using IL-2 IT injections induces no changes in nociceptive thresholds in uninjured naive male and female mice. mTreg expansion in nerve-injured mice induces no changes in (B) acetone, (C) Hargreaves and (D) hotplate behavioral testing. ns = not significant.

**Figure S5. Functional characterization of enkephalin from CD4+ T cells.**

(A) Chimerism of meningeal Tregs and spinal cord microglia in PenkDTR^Δheme^ mice and (B) pooled chimerism comparing Tregs in the meninges and in the lymphoid organs compared to tissue macrophages of the spinal cord (microglia) and the epidermis (Langerhans cells, LC). (C) mTreg number after IP DT injection in PenkDTR^Δheme^ mice. (D) Weight curves of PenkDTR^Δheme^ mice injected with IP DT every three days showing peripheral penk ablation doesn’t induce weight loss. (E) Unaltered spleen weight and (F) meningeal and (G) spleen CD4^+^ T cell populations after IP DT. Unaltered CD4 T cell populations in the (H) spleen and (I) in the nerve and unaltered (J) mouse weight and (K) spleen weight. (L) Graft Versus Host Disease (GVHD) score in *Rag2^-/-^* mice injected with pre-activated Tconv alone or with *Penk^+/+^* or *Penk^-/-^* Tregs, n=3-4 mice per group. (M) IFN- ^+^ and (N) IL-17^+^ CD4^+^ T cells after GVHD induction. ns = not significant, \**p*<0.05, \*\*\**p*<0.001.

