## Supplementary figures and images for "Regulatory T cell-derived enkephalin gates nociception"

### Supplemental Figure 1

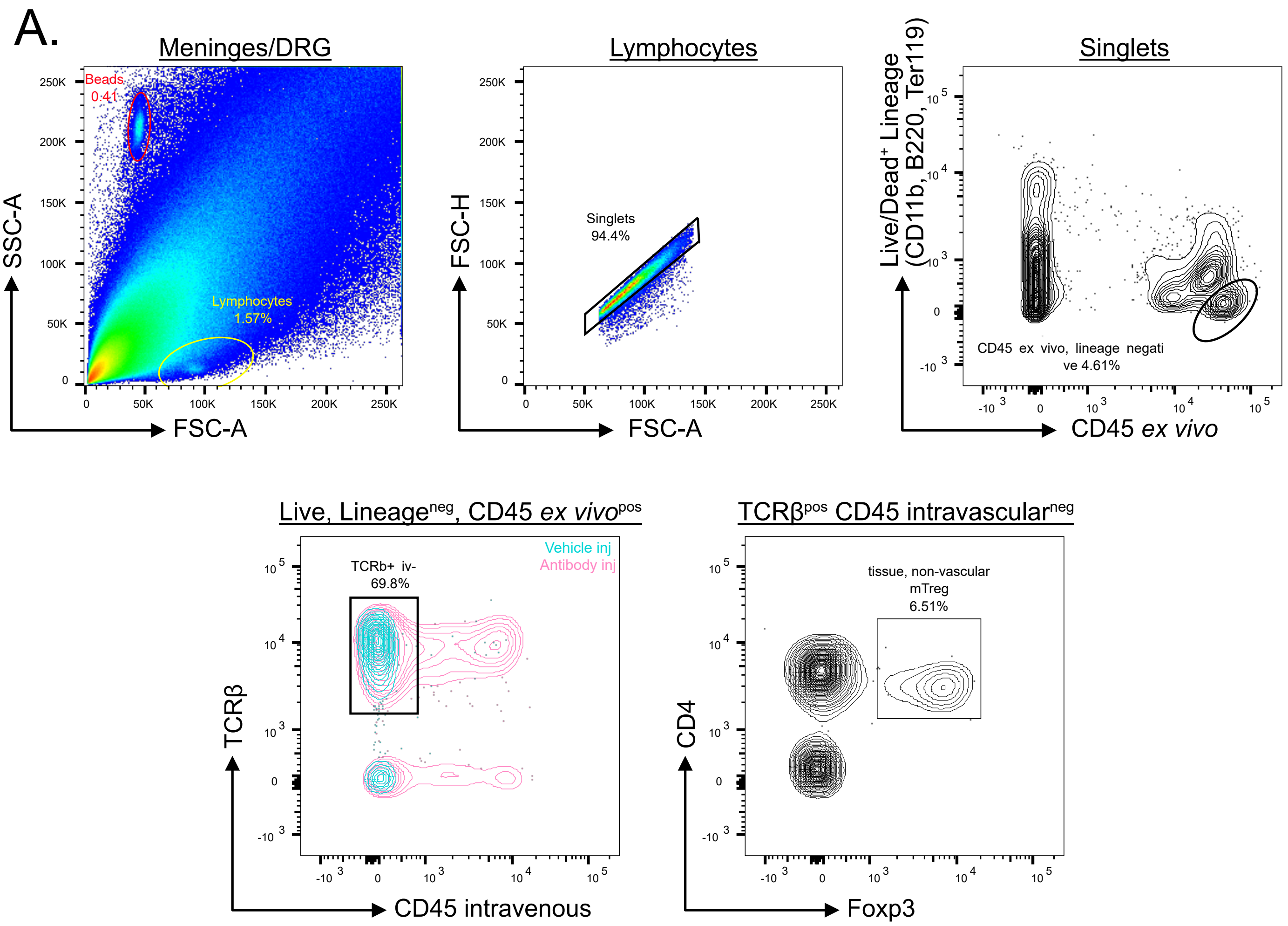

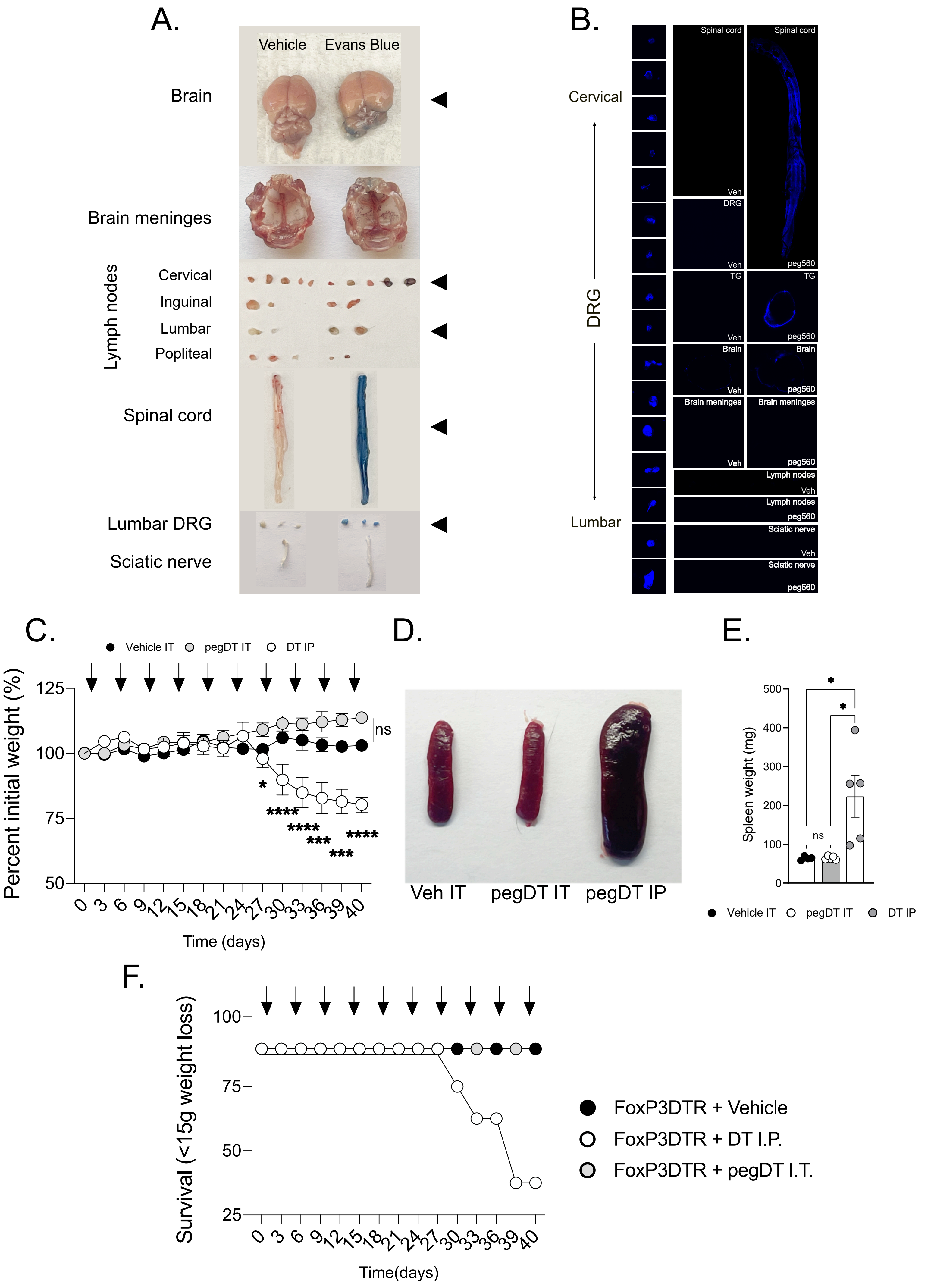

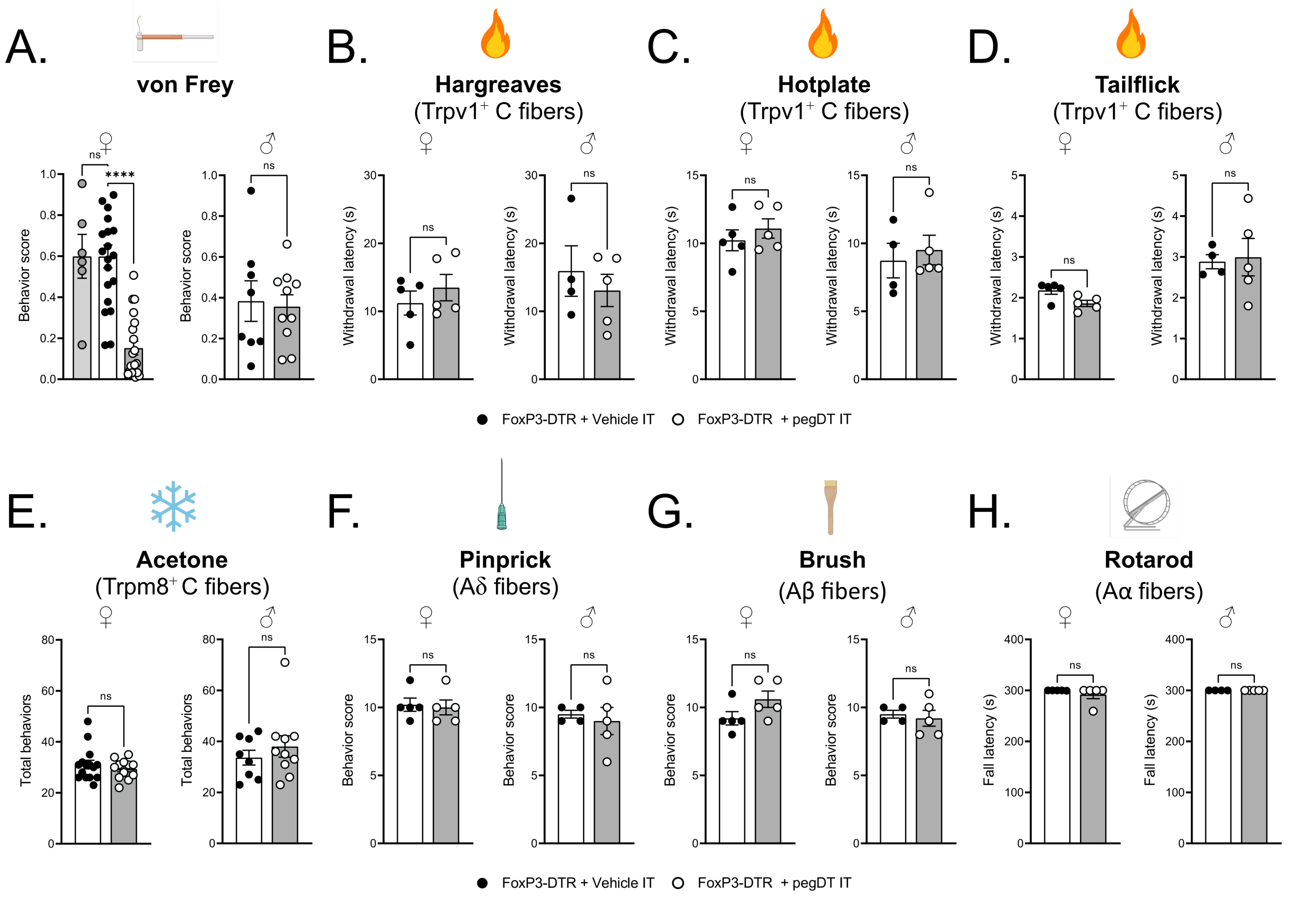

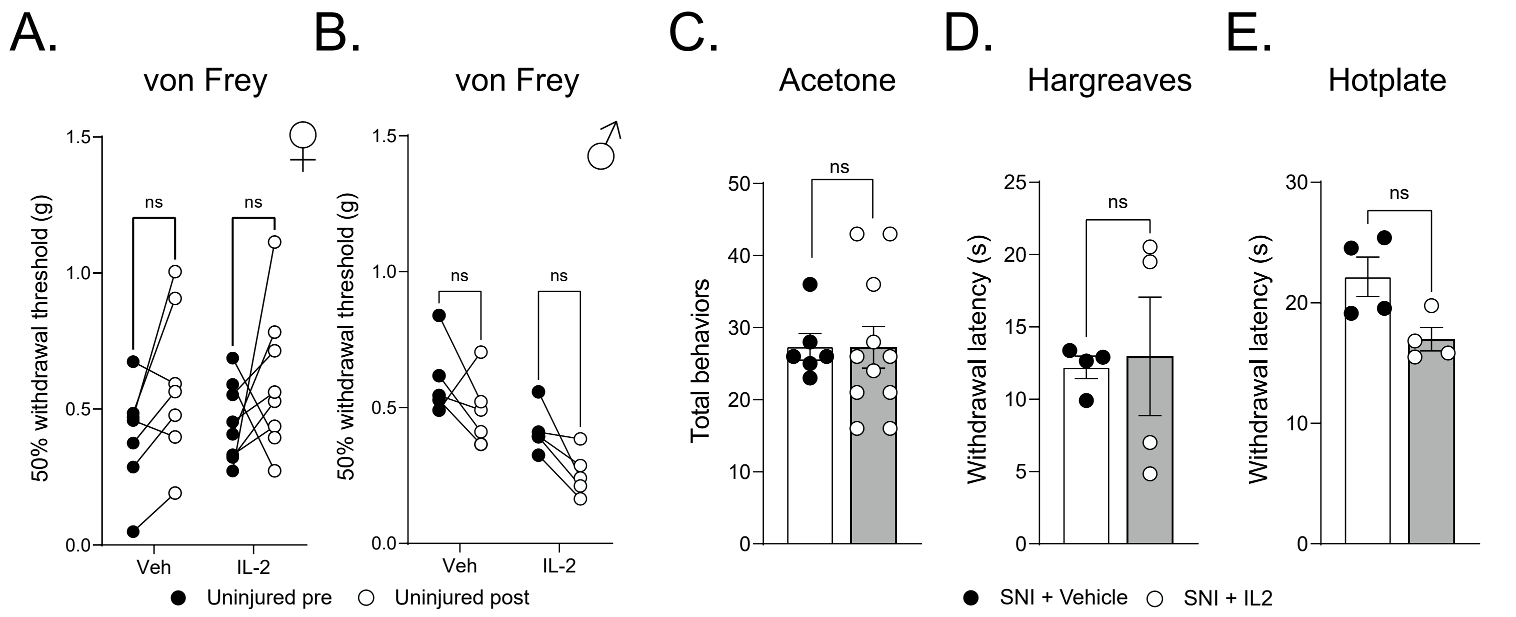

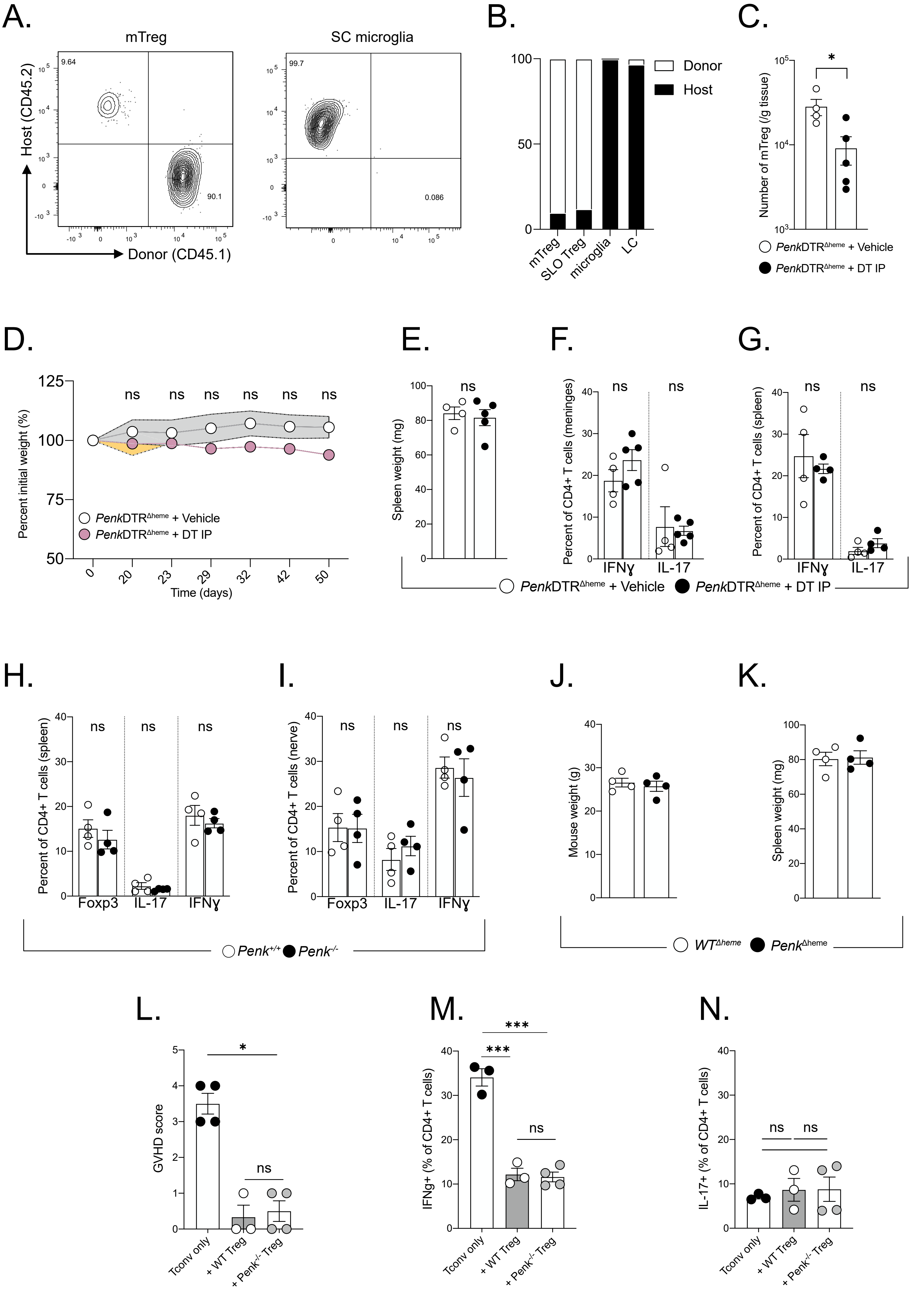

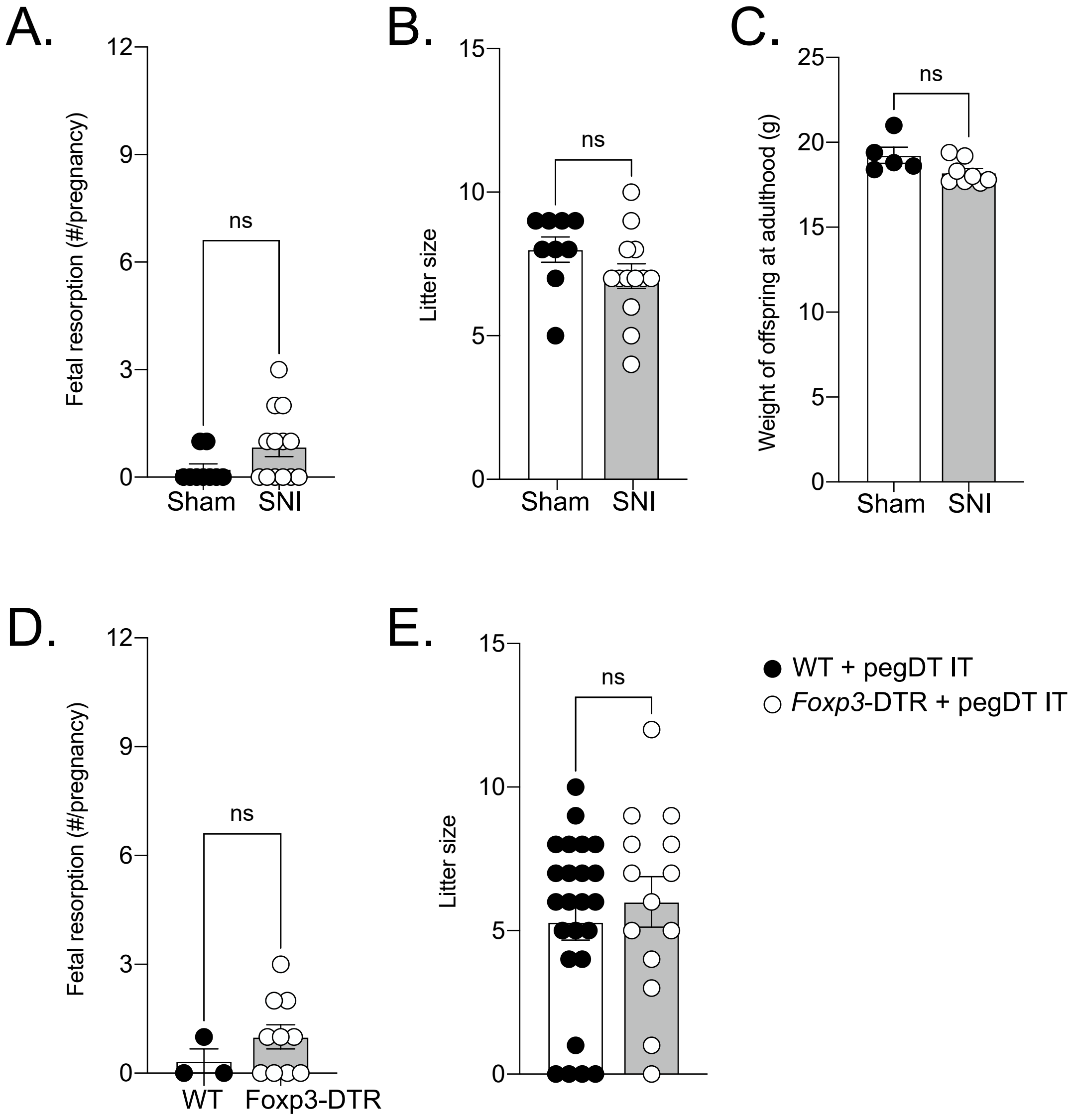

### Supplemental Figure 2

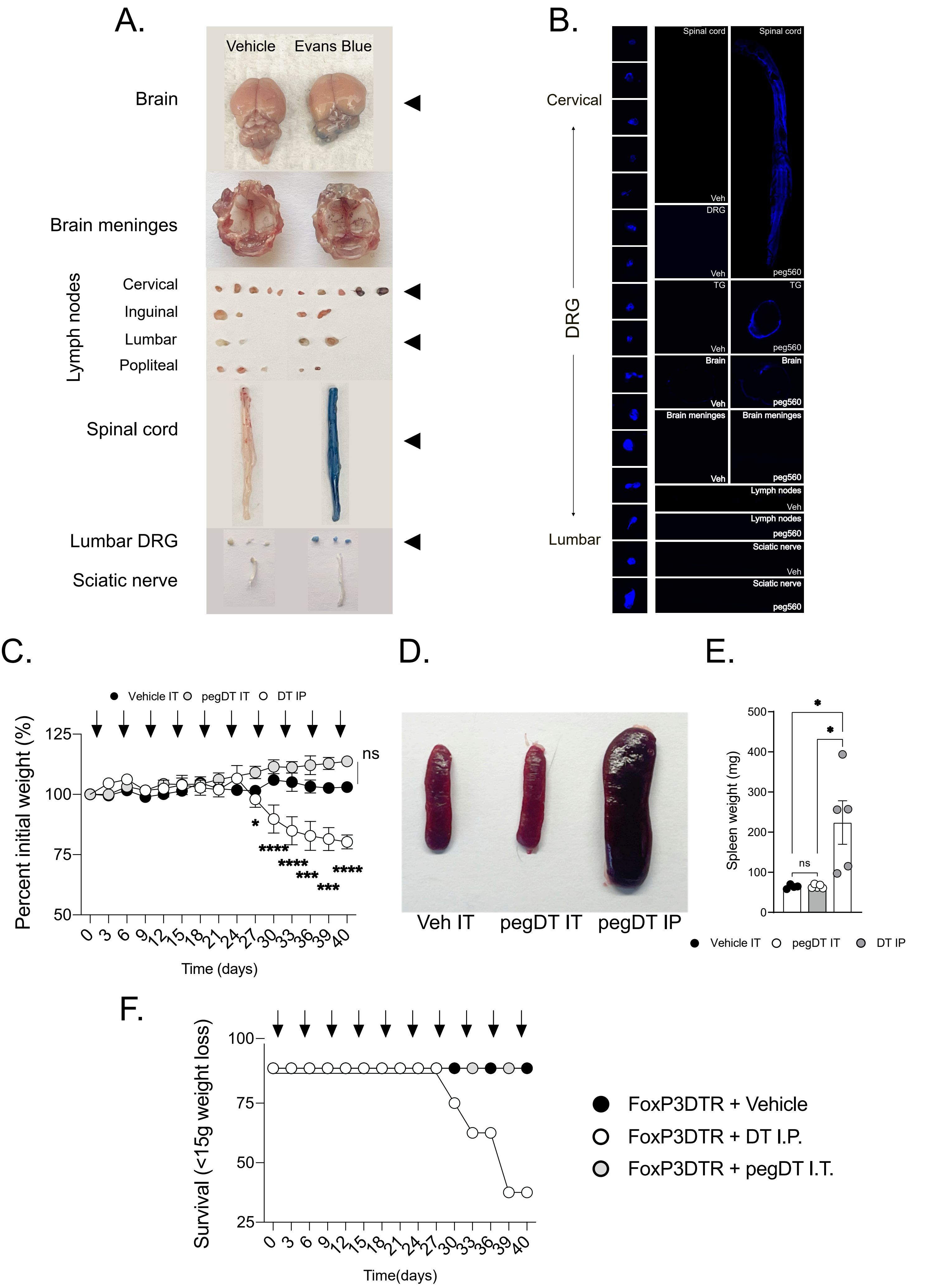

### Supplemental Figure 3

Supplemental Figure S3

Midavaine et al., 2024

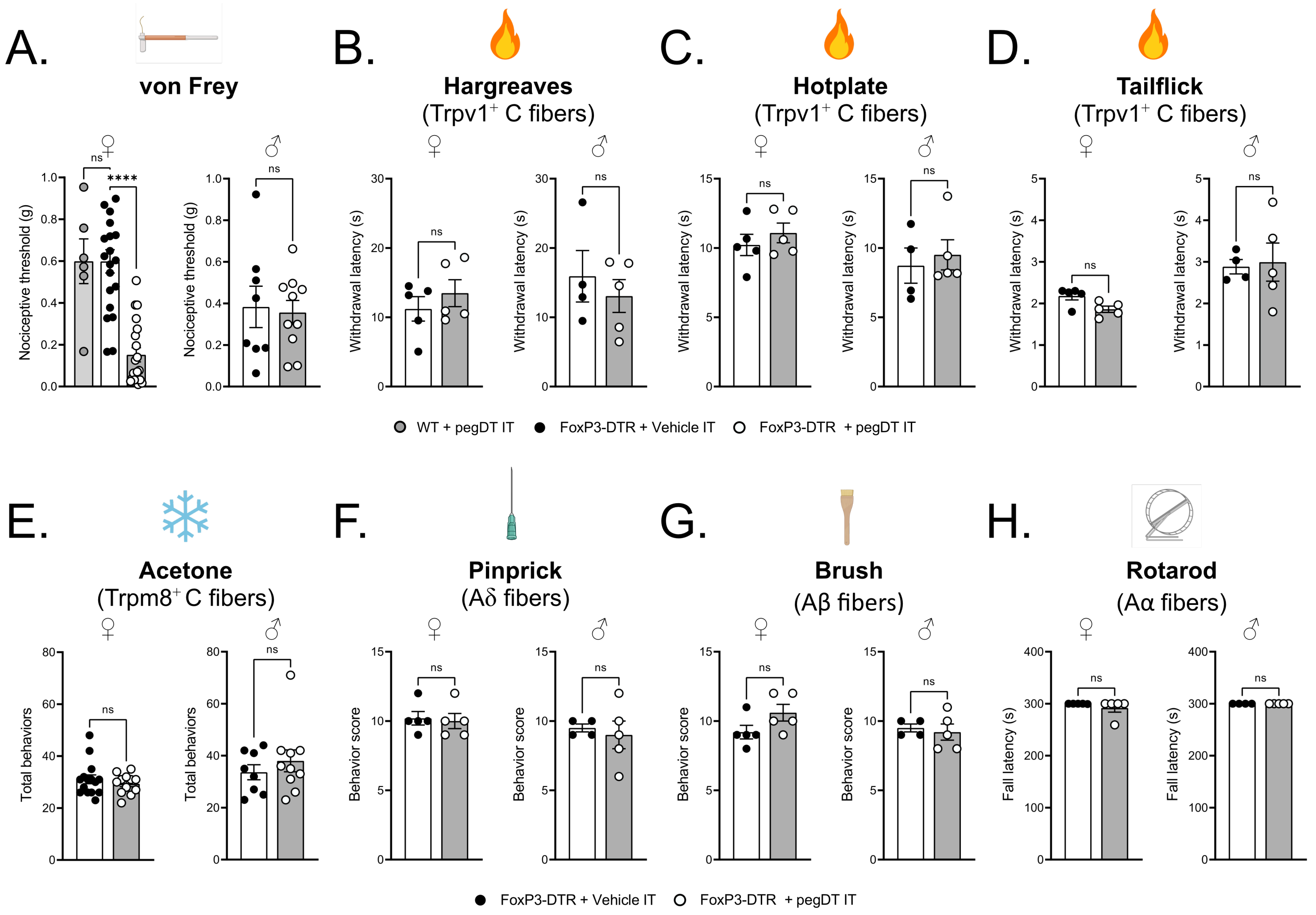

### Supplemental Figure 4

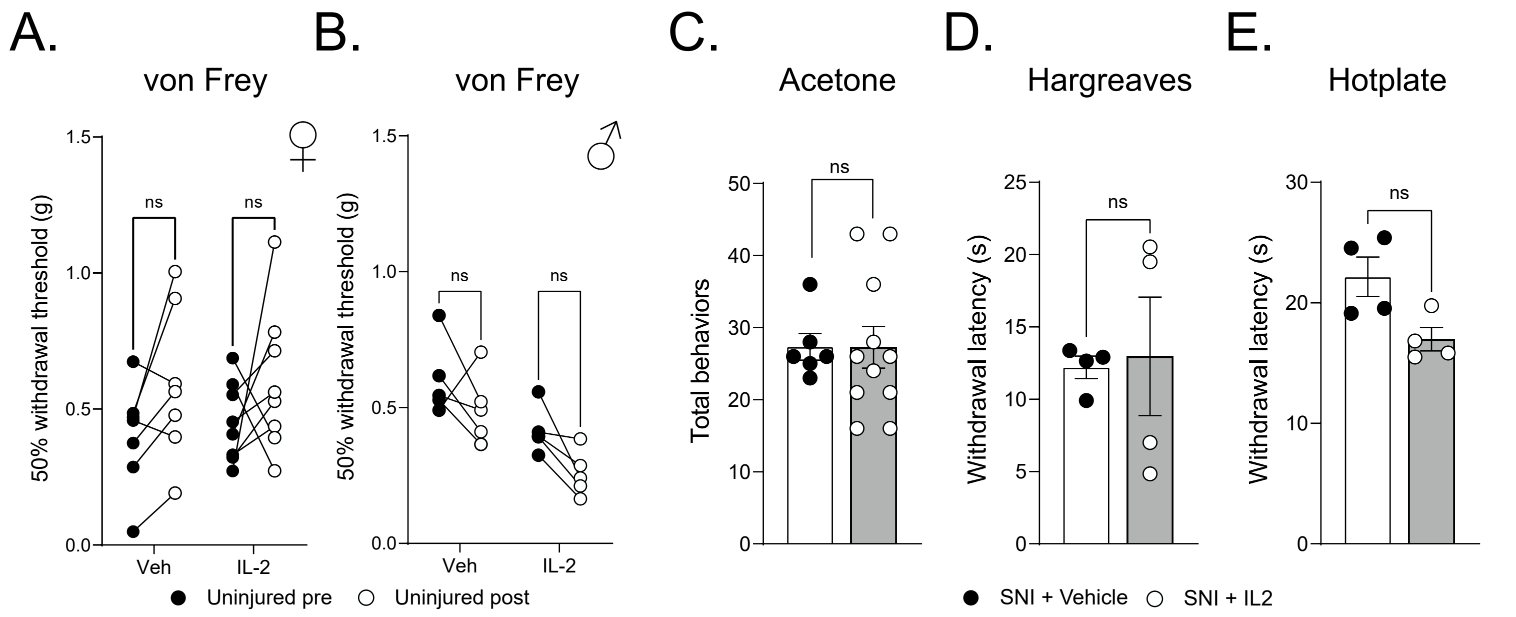

### Supplemental Figure 5

Supplemental Figure S5

Midavaine et al., 2024

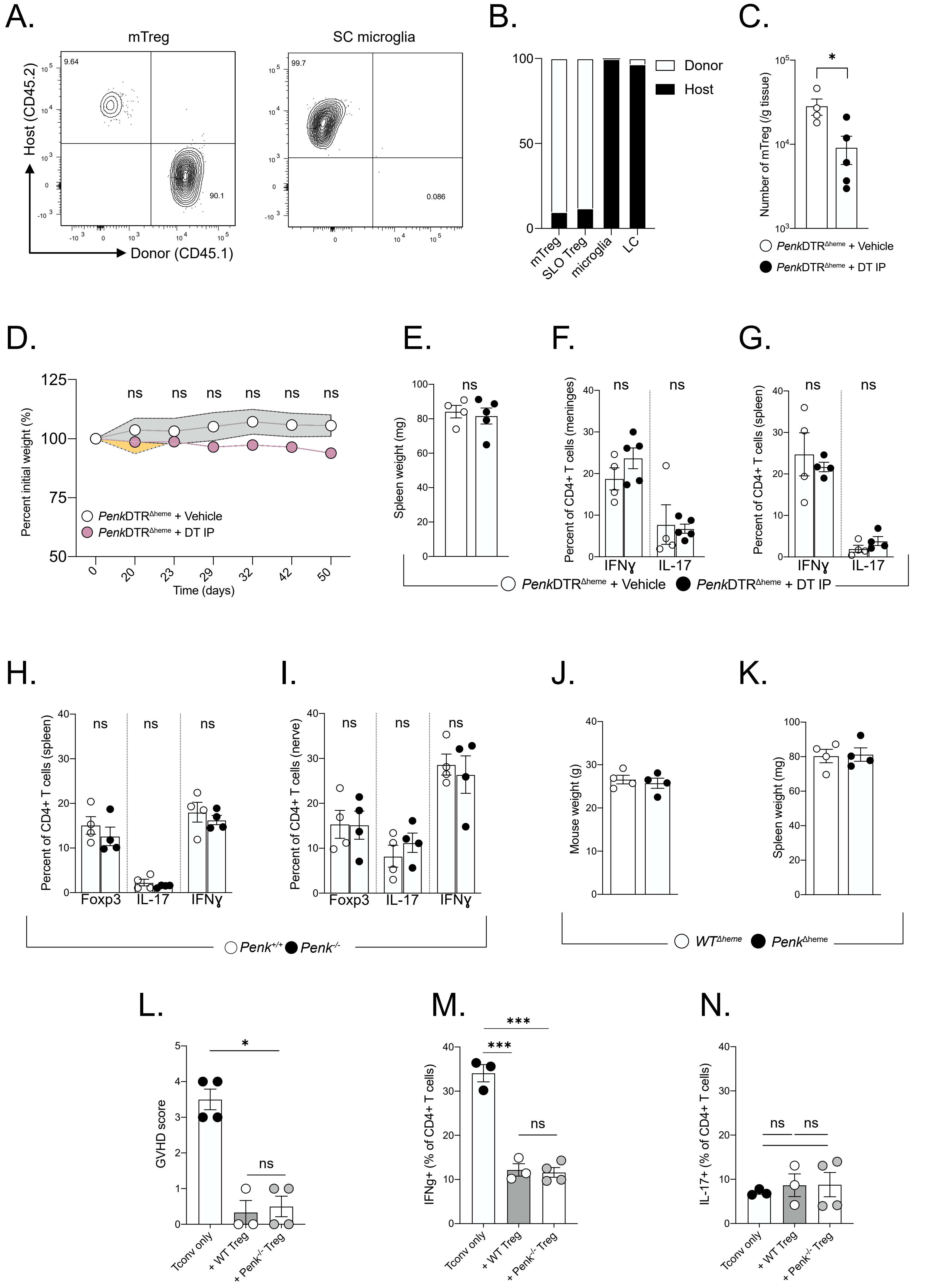
